# High fat diet consumption and social instability stress impair stress adaptation and maternal care in C57Bl/6 mice

**DOI:** 10.1101/2024.05.02.592294

**Authors:** Morgan C Bucknor, Anand Gururajan, Russell C. Dale, Markus J Hofer

## Abstract

Poor maternal diet and psychosocial stress represent two environmental factors that can significantly impact maternal health during pregnancy. While various mouse models have been developed to study the relationship between maternal health and offspring development, few incorporate multiple sources of stress that mirror the complexity of human experiences. Maternal high-fat diet (HF) models in rodents are well-established, whereas maternal psychosocial stress models are still emerging. The social instability stress (SIS) paradigm, serves as a chronic and unpredictable form of social stress. To evaluate the combined effects of a poor maternal diet and social stress on maternal health and behaviour, we developed a novel maternal stress model in adult female C57Bl/6 mice. We observed that all HF+ mice demonstrated rapid weight gain, elevated fasting blood glucose levels and impaired glucose tolerance independent of the presence (+) or absence (-) of SIS. Behavioural testing revealed anxiety-like behaviors remained across all groups prior to pregnancy. However, we did observe a trend of poorer nest quality among all HF+ mice compared to HF-mice following nest building testing. Unlike the other HF+ and HF-stress groups, which exhibited significantly reduced plasma ACTH and corticosterone levels following SIS exposure, we did not observe this reduction in HF+/SIS+ females. In addition, HF+/SIS+ females demonstrated significant postpartum maternal neglect, resulting in fewer numbers of live offspring. These findings suggest that prolonged maternal HF diet consumption, coupled with SIS, places a significant burden on the maternal stress response system, resulting in reduced parental investment and negative postpartum behaviour towards offspring.

## 1. Introduction

Maternal health during pregnancy plays a pivotal role for shaping future health outcomes in offspring (Han et al., 2021a). The intrauterine environment is highly permeable to a range of maternal exposures stemming from the external environment which can significantly alter foetal development (Bronson and Bale, 2016). These environmental factors can be collectively termed as the exposome - some of which include poor maternal diet, sleep, physical exercise (or lack thereof), psychosocial stress, and recurrent pollutant exposure (Han et al., 2021b). Maternal obesity in particular is a growing global concern, given its increasing prevalence and established association with long term metabolic and neurodevelopmental consequences in offspring (Cirulli et al., 2020; Wankhade et al., 2016; Ziauddeen et al., 2022). The CDC reported an 11% increase in pre-pregnancy obesity in the USA from 2016 to 2019, with this trend observed across all age groups (Driscoll and Gregory, 2020). A similar pattern is also becoming evident in the UK (Ziauddeen et al., 2022).

While obesity is multifactorial, the gradual rise in body weight and adiposity is consistent with the current food landscape being dominated by highly palatable foods that are rich in salt, sugar, and fat (Hall, 2018). One of many diet-induced obesity hypotheses suggests that a diet high in fat may be one of the main culprits for this global phenomenon. Dietary fat is more energy dense than protein and carbohydrates and less satiating, leading to increases in overall caloric consumption (Hall, 2018). In young adult male C57Bl/6 mice, researchers modulated different macronutrient compositions between 29 different diets to discover that only increases in dietary fat consumption led to a significant rise in both energy intake and adiposity (Hu et al., 2018). Another C57Bl/6 mouse study in female mice, examined how high fat diet feeding during pregnancy impacts maternal physiology and behaviour, reporting notable increases in cannibalistic episodes of offspring, consistent with reduced neural activity within the olfactory bulbs, a critical brain region for regulating social recognition and maternal behavior (Bellisario et al., 2015, 2014).

Moreover, the impact of stress and the subsequent physiological and psychological response warrants special attention as well. Briefly, the stress response is governed by the hypothalamic-pituitary-adrenocortical (HPA) system which releases glucocorticoids (i.e. cortisol or corticosterone) into the circulation and aids in the mobilisation of energy needed for an organism to cope with adversity (Bartolomucci et al., 2005; Radley and Herman, 2023). However, persistent glucocorticoid secretion can eventually become maladaptive and impair the flexibility of the stress system which challenges successful stress adaptation (Herman, 2013). Chronic, unpredictable stressors, such as social conflict or instability, can place a heavy burden on the HPA system and disrupt the body’s natural ability to return to baseline leading to progressive deterioration in both physiological and behavioural health (Bartolomucci et al., 2005; McEwen and Wingfield, 2003; Ramsay and Woods, 2014).

Preclinical models have been developed in rodents to better understand the impact of chronic social stress exposure on development and behaviour. Commonly used paradigms include chronic social defeat, social isolation and social instability stress (SIS) (Gururajan et al., 2019; Mumtaz et al., 2018; Yohn et al., 2019). The SIS paradigm is a well-validated approach for modelling social stress; especially in female mice as it is one of the few paradigms not reliant on the incidence of aggression. This paradigm continuously disrupts social hierarchies in a group housing environment by changing the cage composition several times per week (Koert et al., 2021a). While there are plenty of iterations of SIS protocols, observations that have been conserved across studies include increases in negative valence behaviours, HPA axis activation and elevations in circulating corticosterone levels (Schmidt et al., 2010a; Yohn et al., 2019). Further, parental exposure to SIS has been shown to directly transmit anxiety and social deficits to subsequent generations of offspring (Saavedra-Rodríguez and Feig, 2013).

In an effort to simulate a real-world context, we developed a ‘two-hit’ environmental stress model which exposed female C57Bl/6 mice simultaneousy to a diet high in fat and 6-weeks of SIS just prior to becoming pregnant. We examined resulting metabolic, hormonal, behavioural and reproductive outcomes following both modes of stress. Our findings demonstrate that the combination of a prolonged high-fat diet and SIS has adverse effects on the maternal stress response and postpartum behaviour.

## 2. Materials and methods

### 2.1 Animals

C57Bl/6 mice were used throughout the study. All female mice (n = 48) were obtained from the Australian BioResources (ABR) facility based in Moss Vale, New South Wales at 5 weeks of age and held at the animal facility at the Charles Perkins Centre (25 °C, 12 h light:dark cycle) with ad libitum water and grain-based meat free chow until commenced on specialised semi purified diets. Each cage consisted of environmental enrichment items and corn cob bedding (Premium Corn Cob Laboratory Animal Bedding, Biological Associates, NSW Australia). Upon arrival, mice were given one week to acclimatise to the holding facility. All animal experiments were performed in compliance with the NSW Animal Research Act and the 8^th^ edition of the NHMRC ‘Australian Code of Practice’ for the care and use of animals for scientific purposes. All efforts were made to minimise animal suffering, to reduce the number of animals used, and to utilise alternatives to in vivo techniques, if available. The study was approved by the University of Sydney Animal Ethics Committee (Protocol 1947).

### 2.2 Diets

Following acclimatisation, female mice were randomly allocated to either a high-fat, high-sugar, no soy semi-pure diet (43% kcal fat diet, SF21-208, Specialty Feeds, WA, Australia) or low-fat, high-sugar, no soy semi-pure control diet formulation (referred to as low-fat diet throughout the study) (12.3% kcal fat diet, SF21-209, Specialty Feeds, WA, Australia) beginning at 6 weeks of age (**Table 1**). The detailed composition of both diets is provided in supplementary file 1. All mice used in the study had ad libitum access to allocated chow and remained on respective diets for the entirety of the study. All mice were weighed once per week for 8 consecutive weeks. Physiological parameters measured included absolute weight gain and percentage (%) of weight gained on a weekly basis.

**Table 1.**
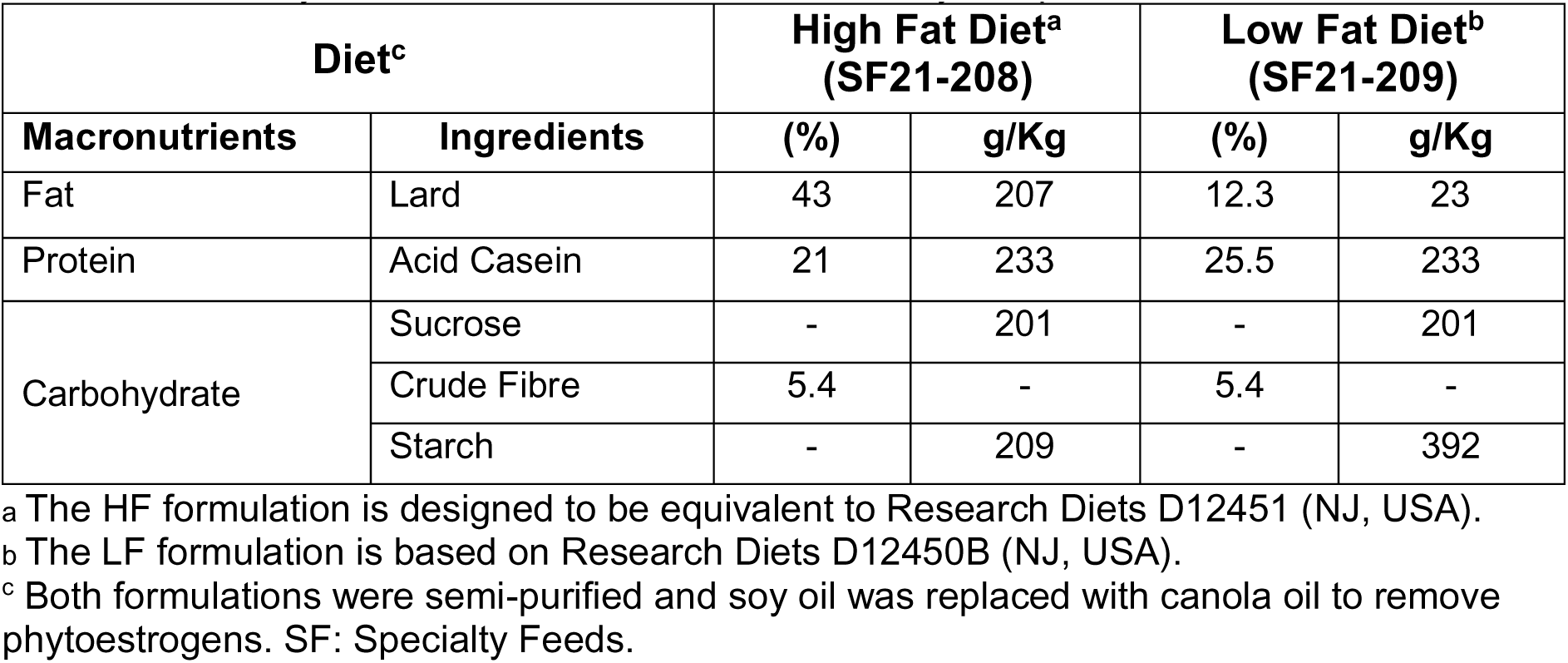
Summary of nutrition facts of HF and LF dietary compositions.

| Diet<sup>c</sup> |  | High Fat Diet<sup>a</sup><br>(SF21-208) |  | Low Fat Diet<sup>b</sup><br>(SF21-209) |  |
| --- | --- | --- | --- | --- | --- |
| Macronutrients | Ingredients | (%) | g/Kg | (%) | g/Kg |
| Fat | Lard | 43 | 207 | 12.3 | 23 |
| Protein | Acid Casein | 21 | 233 | 25.5 | 233 |
| Carbohydrate | Sucrose | - | 201 | - | 201 |
|  | Crude Fibre | 5.4 | - | 5.4 | - |
|  | Starch | - | 209 | - | 392 |

<sup>a</sup> The HF formulation is designed to be equivalent to Research Diets D12451 (NJ, USA).
<sup>b</sup> The LF formulation is based on Research Diets D12450B (NJ, USA).
<sup>c</sup> Both formulations were semi-purified and soy oil was replaced with canola oil to remove phytoestrogens. SF: Specialty Feeds.

### 2.3 Maternal Stress Groups

After randomised stress group allocation the cohort (n = 48) was staggered 2-weeks apart in order to streamline longitudinal experiments. The detailed experimental design can be found in (**Fig. 1A**).

**Figure 1.**
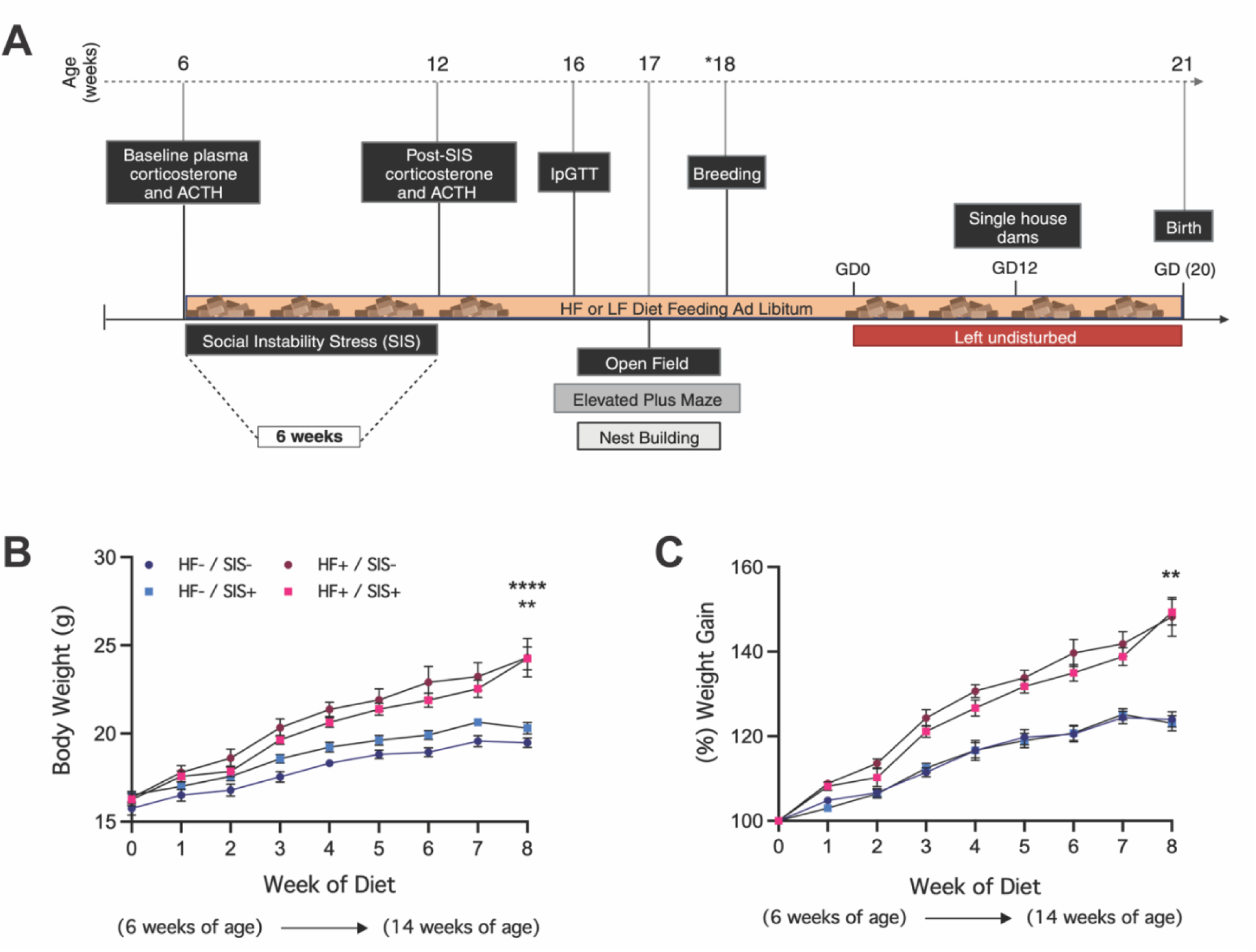
Maternal high-fat diet intake results in rapid weight gain over 8-weeks independent of SIS. (**A**) Schematic of experimental design using BioRender.com. *18 weeks of age = age of female mice during first round of staggered breeding pairings. (**B**) Absolute weight gain across all stress groups. (**C**) Percentage of weight gained over 8-week period. HF- / SIS- (n=8), HF- / SIS+ (n=16), HF+ / SIS- (n=8), HF+ / SIS+ (n=16). Data shown as mean ± SEM. *Denotes statistically significant difference (HF+ / SIS-) and (HF+ / SIS+) vs (HF- / SIS-) and (HF- / SIS+) groups (**p<0.01, **** p<0.0001). Figure B, 2Way RM ANOVA; C, Dunn’s post-hoc test. ACTH = adrenocorticotropic hormone, IpGTT = intraperitoneal glucose tolerance testing. GD = gestational day.

First batch (n = 24):

High-fat diet (HF+) + Social Instability Stress (SIS-) (HF+/SIS-) (n = 4) High-fat diet (HF+) + Social Instability Stress (SIS+) (HF+/SIS+) (n = 16) Low fat-diet (HF-) + Social Instability Stress (SIS-) (HF-/SIS-) (n = 4)

Second batch (n = 24):

High-fat diet (HF+) + Social Instability Stress (SIS-) (HF+/SIS-) (n = 4) Low-fat diet (HF-) + Social Instability Stress (SIS+) (HF-/SIS+) (n = 16) Low fat-diet (HF-) + Social Instability Stress (SIS-) (HF-/SIS-) (n = 4)

### 2.4 Social Instability Stress

The social instability stress (SIS) paradigm has been shown to be effective in inducing psychological stress phenotypes in female rodents with clinical relevance (Koert et al., 2021a; Schmidt et al., 2010a; Yohn et al., 2019). A modified 6-week version of the original protocol was thus used for this study, starting one day after being switched to either the HF or LF diet. Here, group cage compositions (4 mice/cage) were changed at random, twice per week by rotating an individual mouse from each cage into a new, clean cage with 1-3 unfamiliar female cage mates. The rotation schedule was randomised to minimize the frequency of repeated encounters with the same cage mates throughout the experiment. At the end of the SIS exposure, mice remained group housed with cage mates from the last rotation for the remainder of the study.

### 2.5 Plasma Collection

Non-terminal blood collection was done at two timepoints: 1-week after arrival to the animal facility (baseline) and following 6-weeks of SIS (post-SIS); non-SIS mice were included as controls and bled at the same time points. Blood collection took place between 09:30h-12:00h in the morning under mild physical restraint. Mice were habituated for two-consecutive days inside a small transparent cylinder with air holes that served as a restraining device during blood draws on collection day and the procedure room. All blood samples were kept in a 10% volume of heparin (10 Ku/mL) (Sigma, Cat #H3393) on ice and later centrifuged for 10min at 2000xg at 4°C. Plasma was transferred to clean, labelled 1.5ml Eppendorf tube and stored frozen at -80C until determination of corticosterone and adrenocorticotropic (ACTH) hormone concentrations.

### 2.6 Glucose Tolerance

After 8-weeks on respective diets, intraperitoneal glucose tolerance tests (ipGTTs) were performed on all SIS-mice and a subset of SIS+ mice within the cohort at 16-weeks of age. Animals were fasted in the morning for 6-hours prior to testing in the afternoon. Mice were brought into the procedure room and habituated up to 15 minutes prior to the procedure. Sterile glucose (1g/kg body weight) was administered via intraperitoneal (i.p.) injection. Tail vein blood was sampled at 0-, 15-, 30-, 60- and 120-mins as described above (see 2.5). Blood glucose level was measured using Caresens™ N glucometer (i-SENS Inc, Korea). The incremental area under the curve (AUC mmol/L/min) was calculated using mean values per stress group.

### 2.7 Corticosterone and ACTH ELISAs

Plasma corticosterone (ng/ml) (Ab108821) and adrenocorticotropic hormone (pg/ml) (Ab263880) were quantitatively measured using commercial ELISA kits according to the manufacturer’s instructions. Plasma samples were diluted 1:8 or 1:100 for corticosterone assay and 1:4 or 1:8 for the ACTH assay. All samples were analysed in duplicate. Absorbance was measured at 450nm using a Tecan infinite M1000 Pro plate reader. Concentrations of each protein were calculated using the free online AssayFit Pro 96 well curve fitting ELISA calculator (https://www.assayfit.com/home.html) and standard curves were generated by fitting the data to a 4-parameter logistic curve (4PL).

### 2.8 Behavioural Testing

Behaviour testing was performed before pregnancy, at 17-weeks of age, during week-10 of respective diets and following 6-weeks of SIS stress (if applicable) on all mice. At least 1hr prior to the start of testing, mice were acclimatised in their home cages to a dimly lit empty holding room adjacent to the testing room. The following behavioural battery was performed to examine anxiety-like behaviour and locomotor activity: elevated plus maze, open field, and nest building. The same testing order was kept consistent between batches of females. Elevated plus maze and open field testing were performed in the same testing room between 10:00-14:00hrs during the standard light phase with one rest day in-between tests. Each apparatus was cleaned thoroughly with 80% ethanol solution between test mice. Nest building was performed following the end of open field testing where females were left singly housed, with ad libitum access to food and water, overnight till nest assessments the next morning. Raw data of elevated plus maze and open field tests was analysed using AnyMaze video tracking software (V7.20).

### 2.9 Elevated Plus Maze Test

Mice were placed into the centre of the plus-shaped maze facing the open arm and allowed to freely explore for a 5-minute testing period. The centre of the maze was brightly lit (250-400 lux) to be a truly potent stimulus that can best assess anxiety-like behaviour. The maze was positioned in the centre of the testing room, 43 cm off the floor. Each arm was 30 cm in length, 5.5 cm in width and the height of the black plexiglass wall surrounding the closed arms was 16 cm. Parameters measured included total distance travelled, percentage (%) of open arm entries and the percentage (%) of time spent in the open arms.

### 2.10 Open Field Test

A mouse was placed into a novel open field arena measuring 40 cm L x 40 cm W and 20 cm in height with a dark plexiglass wall. The centre of the arena was dimly lit (<50 lux) to encourage exploration and the mouse was allowed to freely explore for a 10-minute testing period. Parameters measured included total distance travelled, number of entries to the centre zone and the percentage (%) of time spent in the centre zone.

### 2.11 Nest Building

Nest building is an innate behaviour in male and female rodents and nesting material is highly valued by mice. Typically, changes in nesting behaviour are indicative of a change in health or welfare and can be sensitive to physiological and/or environmental manipulations (Neely et al., 2019). Therefore, in this study we used an existing protocol (Deacon, 2006) to test whether or not the compounding effects of HF diet or SIS would alter nesting behaviours.

Briefly, mice were singly housed overnight, with ad libitum access to food and water in standard cages (36 cm L x 18 cm W x 13 cm H) containing paper-based purocel bedding (Pura Paper Premium (Cellulose bedding), Able Scientific, WA Australia) and a single cotton Nestlet square. No additional environmental enrichment items were provided. The next morning, nests were assessed on a rating scale of 1-5 as follows: (1) nestlet not noticeably touched, (2) nestlet partially torn, (3) nestlet mostly shredded but often no identifiable nest site, (4) an identifiable but flat nest and (5) a near perfect nest.

### 2.12 Behavioural z-score normalisation

Behavioural z-score normalisation was calculated using the following output from each behavioural test: **open field**: time in the centre zone and total distance travelled, **elevated plus maze**: time in the open arms and total distance travelled, **nest building**: the percentage of nestlet material used.

The following formula was used to calculate each individual z-score per test (Kraeuter, 2023):

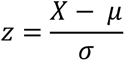

X = observed output

μ = group mean

σ = population standard deviation

The integrated anxiety z-score was calculated by taking the sum of each individual z-score from each test and dividing by the number of tests included in the analysis:

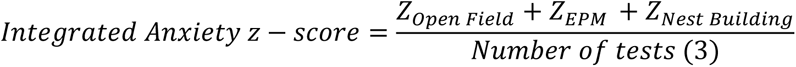

### 2.13 Statistical Analysis

GraphPad Prism (10.1.0) (GraphPad Software Inc., La Jolla CA, USA) and RStudio (V2023.09.1+494) were used for statistical analyses. All data were checked for normality and homogeneity of variance using Shapiro Wilk and Bartlett’s tests, respectively. Where applicable, parametric (RM 2Way ANOVA and 2Way ANOVA) or non-parametric (Friedman, Kruskal-Wallis test) testing was performed. If < 50% of samples within a dataset were not normally distributed, parametric testing was performed. If ≥ 50% samples within a dataset were not normally distributed, non-parametric testing was performed. Multiple comparison post-hoc tests were used where ANOVA, Friedman or Kruskal-Wallis testing suggested statistical significance (p<0.05). All statistical reports are presented in supplementary statistical reports 1-6.

## 3. Results

### 3.1 HF+ mice show rapid body weight gain independent of additional SIS exposure

Long-term high-fat diet consumption results in rapid fat accumulation and subsequently, increased body weight in all female mice. To determine whether the addition of SIS impacted body weight gain during the 6-week stress protocol, all mice were weighed weekly for 8 weeks (**Fig. 1A-C**). Firstly, notable increases in weight gain were detected in all HF+ mice (HF+/SIS- and HF+/SIS+) beginning as early as week 3 of feeding (+1.94g avg mean difference vs LF groups) (**Suppl. Table 1**). 2-Way ANOVA testing revealed a significant interaction between time on the diet x stress group (ANOVA; (F(24, 352) = 5.436, p <0.0001)). By 8 weeks of ad libitum access, both HF+ groups were overweight compared to both LF groups – irrespective of SIS status (+4.37g avg mean difference) (**Fig. 1B, C**). Therefore, concurrent SIS does not significantly influence weight gain.

### 3.2 HF+/SIS- and HF+/SIS+ mice display elevated fasting blood glucose and impaired ipGTT glucose response after 8-weeks of ad libitum consumption

To assess to what degree the HF diet may be altering maternal metabolism, we performed intraperitoneal glucose tolerance tests on all control mice (HF-/SIS-, HF+/SIS-, n = 8/group) and a random subset of mice from each SIS group (n = 8/group) after 8-weeks of diet exposure and 6-weeks of SIS (**Fig. 1A**). Fasting blood glucose levels of all HF mice were increased compared to both LF groups following a 6h fast (+11 mmol/L avg mean difference). However, only HF+/SIS-mice were found to be significantly elevated compared to HF-/SIS-mice (+13.75 mmol/L mean difference, p = 0.0201; Dunn’s post-hoc test) (**Fig. 2A**). Following i.p. administration of a glucose bolus (1 g/kg body weight), both HF groups demonstrated elevated ipGTT response curves compared to both LF groups independent of SIS (**Fig. 2B**).

**Figure 2.**
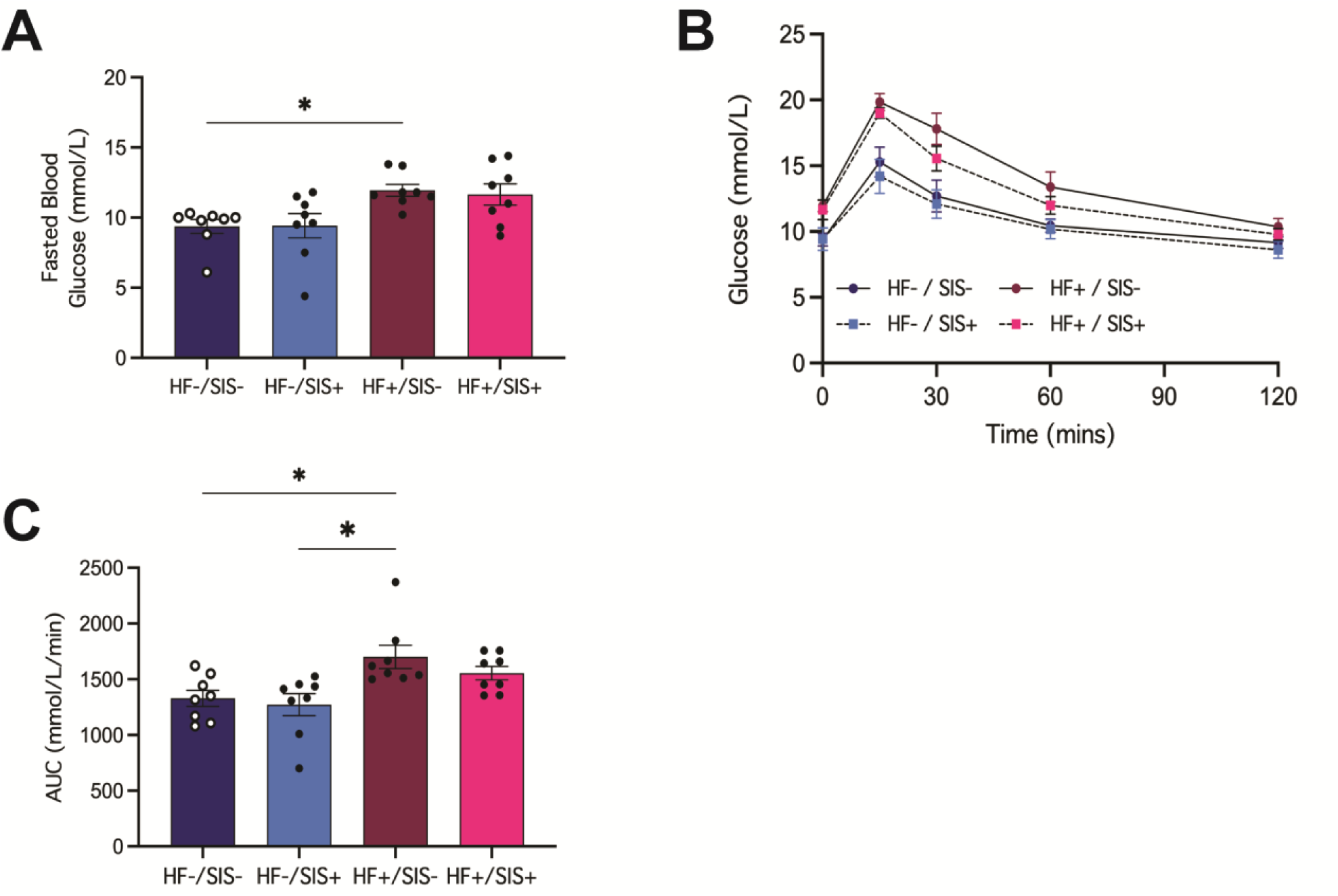
Glucose tolerance is significantly impaired between HF+/SIS- and both LF maternal stress groups. (**A**) Fasted blood glucose in mmol/L following a 6-hr morning fast. (**B**) Mean glucose in mmol/L during ipGTTs is shown following 15-, 30-, 60- and 120- mins post-glucose administration at 16 weeks of age. (**C**) AUC is shown as mmol/L/min as quantification of glucose response during the test period. HF- / SIS- (n=8), HF- / SIS+ (n=8), HF+ / SIS- (n=8), HF+ / SIS+ (n=8). Data shown as mean ± SEM. *Denotes statistically significant difference (HF+ / SIS-) vs (HF- / SIS-) and (HF- / SIS+) (*p<0.05). Figure B-C, Dunn’s post-hoc test.

Yet, the AUC of HF+/SIS- mice was the sole group found to be statistically significant compared to both LF groups (versus (HF-/SIS+) +14.69 mmol/L/min mean difference, p = 0.0104, versus (HF-/SIS-) +13.38 mmol/L/min mean difference, p = 0.0261; Dunn’s post-hoc test) (**Fig. 2C**). Overall, both HF groups demonstrate impaired glucose homeostasis compared to LF groups following 8-weeks of diet intake, indicating the beginning of a metabolic shift towards metabolic syndrome.

### 3.3 HF+/SIS+ mice do not show significant reductions in plasma ACTH and corticosterone following SIS

To determine whether SIS exerted any affect on the neuroendocrine system, we measured levels of two primary stress-related hormones, ACTH and corticosterone at two time points: baseline and post-SIS and/or 6-weeks of respective dietary intake.

Plasma ACTH was found to be significantly altered by time (ANOVA; F(1, 36) = 15.87, p = 0.0003). Although we did not find a significant interaction between stress group x time, Fisher’s post-hoc test revealed significantly decreased levels from baseline – post SIS amongst both LF diet groups independent of SIS status (HF-/SIS- : 109.5 pg/ml mean difference, p = 0.035) ; HF-/SIS+ : 121.3 pg/ml mean difference, p = 0.005) (**Fig. 3A**). While, both HF diet groups did not reveal statistically significant changes post SIS (HF+/SIS- : 96.81 pg/ml mean reduction, p = 0.06 ; HF+/SIS+ : 35.99 pg/ml mean reduction, p = 0.383). However, it is worth noting that the HF+/SIS-group were closer to the verge of statistical significance than HF+/SIS+ mice (**Fig. 3A**). Fisher’s test also reflected that post SIS ACTH levels were much higher on average in HF+/SIS+ mice compared with all other stress groups (HF-/SIS- vs HF+/SIS+ : -125.5 pg/ml mean reduction, p = 0.016; HF-/SIS+ vs HF+/SIS+ : -108 pg/ml mean reduction, p = 0.02; HF+/SIS- vs HF+/SIS+ : -116.3 pg/ml mean reduction, p = 0.026) (**Fig. 3A**).

**Figure 3.**
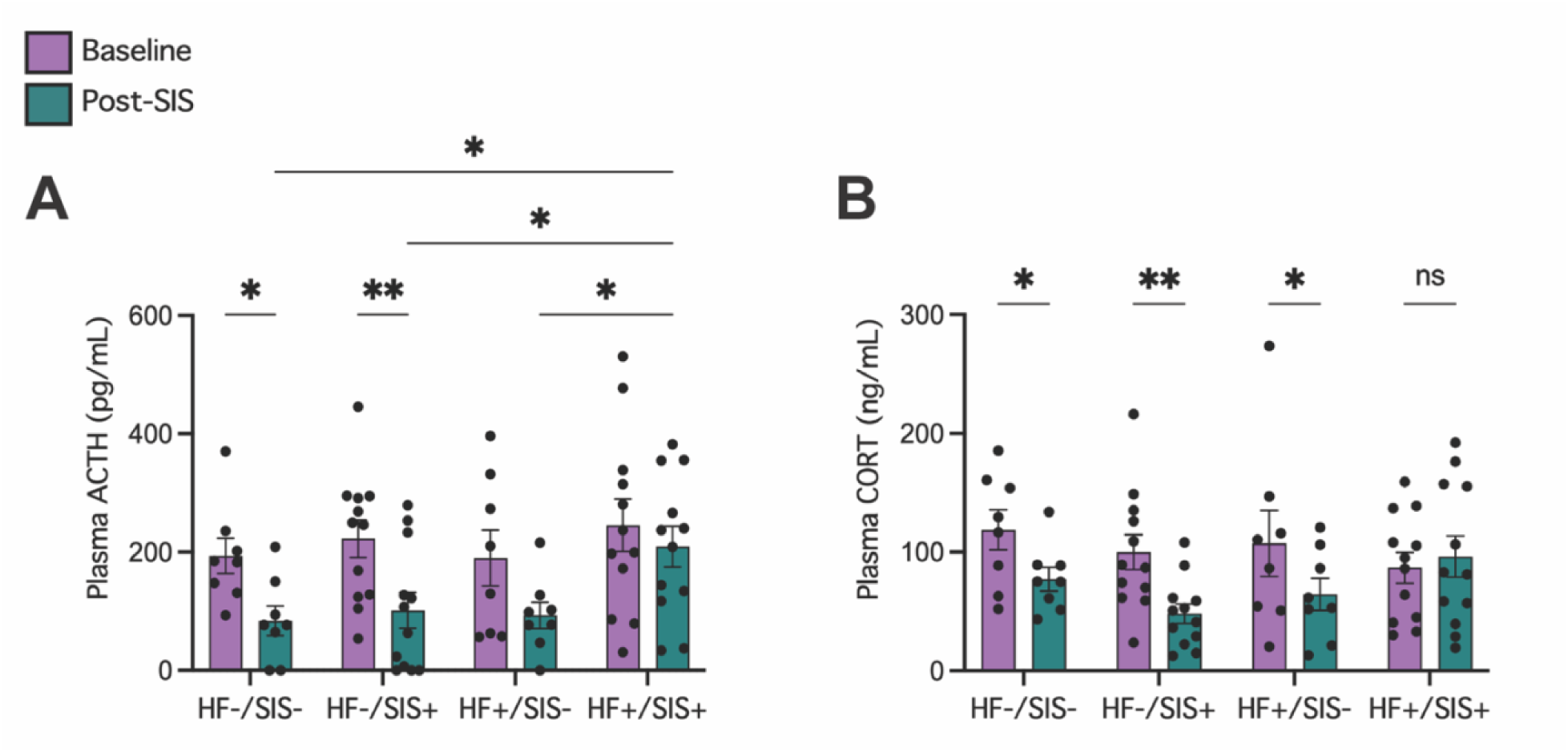
HF+/SIS+ mice do not display significant reductions in plasma ACTH and corticosterone post-SIS. **(A)** Plasma adrenocorticotropic hormone (pg/mL) concentrations measured at baseline and post-SIS stress. HF- / SIS- (n=8), HF- / SIS+ (n=12), HF+ / SIS- (n=8), HF+ / SIS+ (n=12). **(B)** Plasma corticosterone (ng/mL) concentrations measured at baseline (6 weeks of age) and post-SIS stress (12 weeks of age) HF- / SIS- (n=8), HF- / SIS+ (n=12), HF+ / SIS- (n=8), HF+ / SIS+ (n=12). Data shown as mean ± SEM. (*p<0.05, **p<0.01). Figure A-B, 2Way RM ANOVA, A: Fisher’s post-hoc test, B: Tukey’s post-hoc test.

Two-way RM ANOVA revealed a significant interaction between stress group x time (ANOVA; F(3, 36) = 3.059, p = 0.0405) in regard to plasma corticosterone levels (ng/ml). However, time had the largest affect on corticosterone post SIS (ANOVA; F(1, 36) = 13.29, p = 0.0008) (**Fig. 3B**). On average, plasma corticosterone levels were found to be significantly reduced following SIS compared to basal concentrations amongst all stress groups - with the exception of HF+/SIS+ mice (**Fig. 3B**). Tukey’s post-hoc test revealed the HF+/SIS+ group did not demonstrate a statistically significant mean difference in plasma corticosterone between baseline and post SIS (-9.346 ng/ml mean difference, p = 0.5523). Whereas, remaining stress groups did display a large mean difference following SIS (HF-/SIS- : 41.47 ng/ml, p = 0.036; HF-/SIS+ : 51.95 ng/ml, p = 0.002; HF+/SIS- : 42.95 ng/ml, p = 0.030) (**Fig. 3B**). Overall, these results indicate that HF+/SIS+ mice do not experience significant stress-related adaptations following 6-weeks of SIS and HF diet intake.

### 3.4 Open field behaviours remain unchanged across all maternal stress groups

In the open field test, we did not find a significant interaction between the total distance travelled, number of entries to the centre zone or percentage of time spent in the centre zone x all stress groups (**Fig. 4A-C**). This finding is consistent with our pilot data showing the same non-significant trend during open field testing (**Suppl. Fig. 2**).

**Figure 4.**
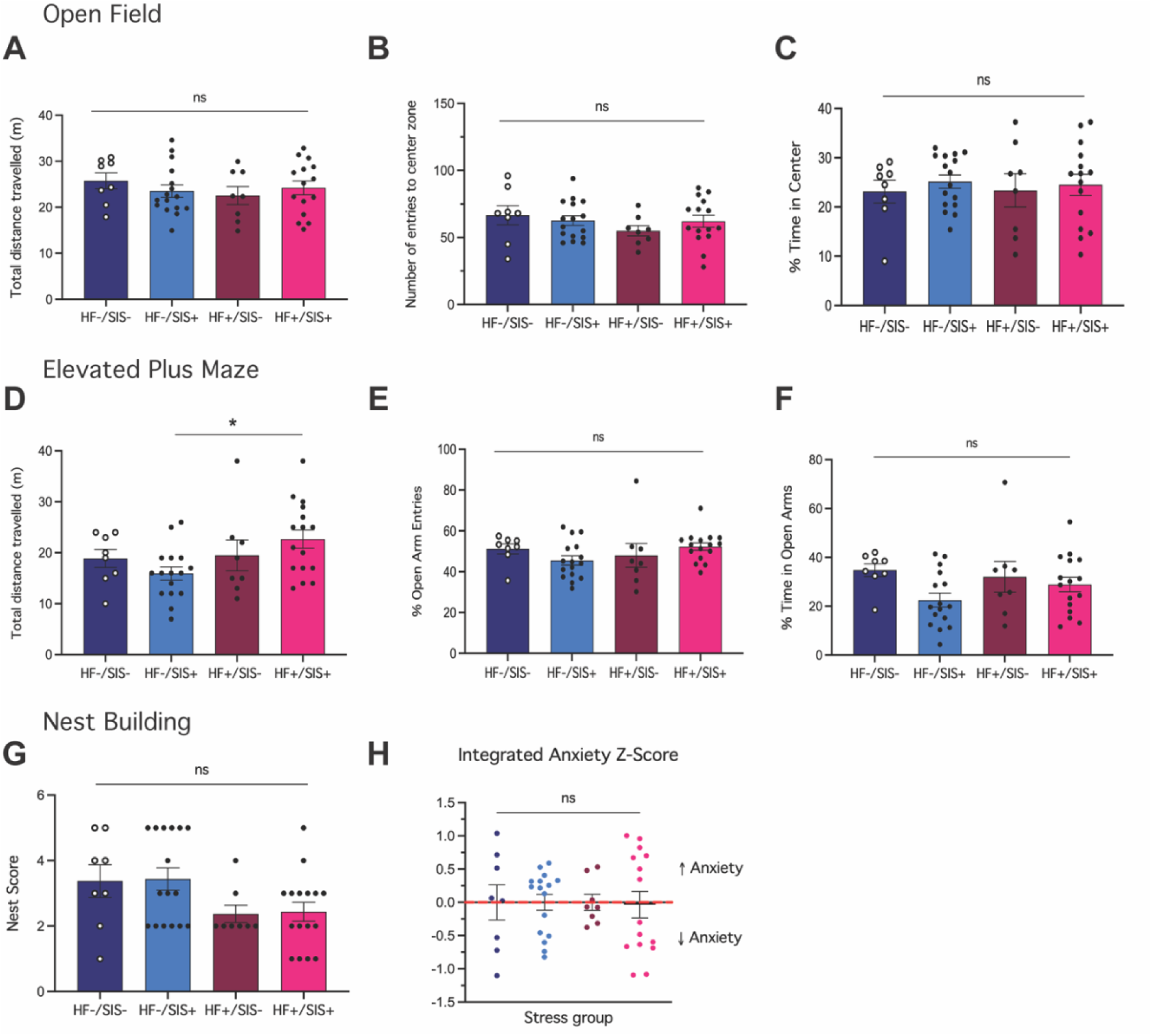
Maternal HF dietary exposure and/or SIS do not significantly impact anxiety- like behaviours prior to pregnancy. (**A**) Total distance travelled (m) during the open field test, (**B**) number of entries to the center zone and (**C**) percentage of time spent in the center zone. HF-/SIS- (n=8), HF-/SIS+ (n=16), HF+/SIS- (n=8), HF+/SIS+ (n=15). (**D**) Total distance travelled (m) during the elevated plus maze test, (**E**) percentage of open arm entries and (**F**) percentage of time spent in the open arms. (**G**) Nest score following 24hr testing period.) (**H**) Integrated anxiety z-score of all behavioural measures. Tests performed at 17 weeks of age. HF-/SIS- (n=8), HF-/SIS+ (n=16), HF+/SIS- (n=8), HF+/SIS+ (n=16) for elevated plus maze, nest building and Z-score. Data shown as means ± SEM. (*p<0.05), ns = not significant. Figure A, D-F, Sidak’s post-hoc test, B-C, Tukey’s post-hoc test, G-H, Dunn’s post-hoc test.

### 3.5 HF+/SIS+ mice demonstrate increased locomotor activity during elevated plus maze

During the elevated plus maze test there was no significant interaction found between stress group x the percentage of entries and percentage of time spent in the open arms (**Fig. 4E, F**). However, there was a marginal main effect observed between stress groups in the context of the total distance travelled during the test (ANOVA; F(3, 29) = 2.873, p = 0.0533). Sidak’s post-hoc test revealed HF+/SIS+ mice travelled a farther distance on average during the test compared to HF-/SIS+ mice, (+6.75m mean difference, p = 0.0389) (**Fig. 4D**). Therefore, while all mice behaved similarly across stress groups, the HF+/SIS+ mice generally demonstrated increased locomotor activity.

### 3.6 Nest building behaviours are not statistically impacted by diet or SIS prior to conception and confirmed by integrated anxiety Z-scores

Following the elevated plus maze and open field tests, all mice were single housed overnight, with access to food and water, and left with a cotton nestlet square to construct nests in the absence additional housing enrichment. Nest quality was scored on a scale of 1-5 (see 2.11). While there was a downward trend observed in overall nest quality (i.e., HF+ animals constructing lower quality nests), we did not detect any statistically relevant changes between all maternal stress groups (p = 0.0913; Kruskal-Wallis) (**Fig. 4G**).

To gather a more complete summary of how anxiety was impacted following maternal high fat diet intake (+/-) and SIS (+/-), we calculated an integrated z-score statistic of all behavioural tests described above (see 2.12). This summary z-statistic confirmed that anxiety-like behaviours remained largely unaffected by HF diet and/or SIS (**Fig. 4H**).

### 3.7 HF+/SIS+ and HF+/SIS-dams display frequent episodes of maternal neglect towards offspring

Following metabolic, hormonal, and behavioural characterisation of the maternal stress model our final objective was to assess how HF diet and/or SIS impacted fertility, maternal care, and importantly, neglect. Maternal care is defined here as feeding and nurturing offspring, while neglect we defined here as litter cannibalisation or pups found dead shortly after birth. Of the 48 total females from the cohort, 45 were paired for breeding. This was due to resourcing constraints and a sufficient number of offspring having been generated from this group. All 45 breeders did not demonstrate any alterations to fertility as they were all able to conceive and carry pups full-term. We observed significant proportions of mice from both HF groups to exhibit the most maternal neglect as opposed to both LF groups; with both LF groups exhibiting the most maternal care (Chi-square test: X^2^(3, N = 45) = 65.69, p<0.0001) **Suppl. Table 2**, **Fig. 5A**). Of the dams that demonstrated maternal care, litter sizes were largely consistent across all stress groups and weanling weight was modestly higher from HF dams opposed to LF dams (**Suppl. Table 1**). Moreover, we observed a significant trend with a larger proportion of females from HF+/SIS+ dams and larger proportion of males from HF+/SIS-dams compared to LF dams where the male:female sex distribution was closer to 50:50 (Chi-square test: X^2^(3, N = 113) = 37.10, p<0.0001) (**Suppl. Table 2, Fig. 5B**).

**Figure 5.**
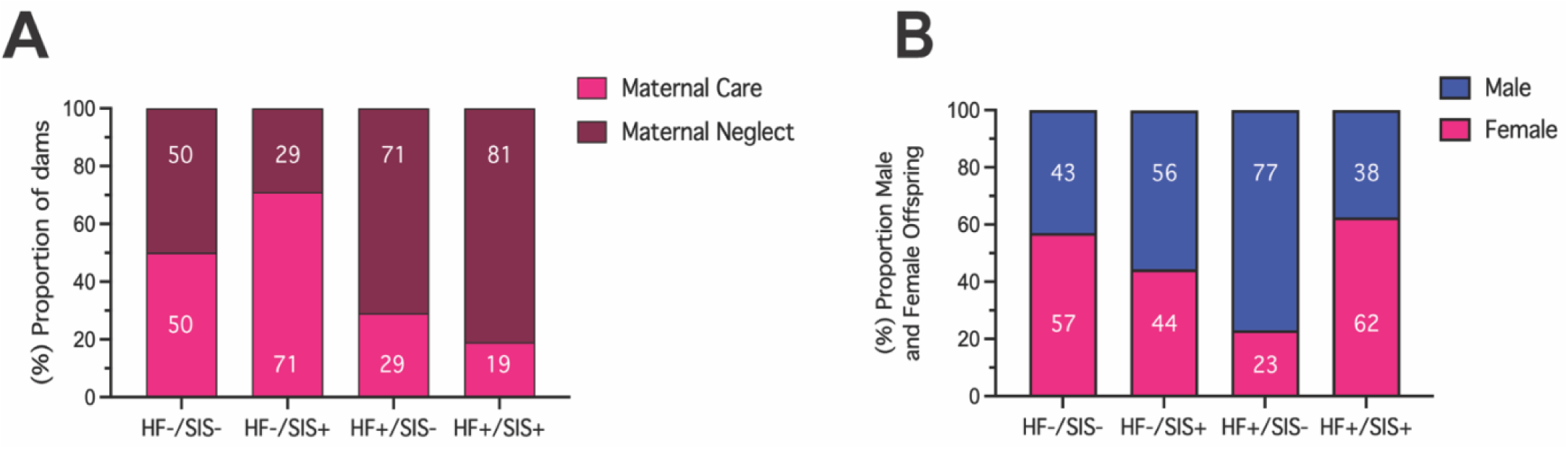
HF+/SIS- and HF+/SIS+ mice display more frequent postpartum maternal neglect and produce skewed sex distribution within litters. (**A**) Percentage of dams exhibiting maternal care or neglect across stress groups. HF-/SIS- (n=8), HF-/SIS+ (n=14), HF+/SIS- (n=7), HF+/SIS+ (n=16). (**B**) Percentage of live male and female offspring across maternal stress groups. Total number of live offspring from each group: HF-/SIS- (n=30), HF-/SIS+ (n=54), HF+/SIS- (n=13), HF+/SIS+ (n=16). Data shown as absolute proportions. Refer to Supplementary table 2 for relevant statistical results.

These findings indicate that HF diet alone, and in combination with SIS results in a higher frequency of maternal neglect toward offspring. Of the HF dams that did care for offspring (5 out of 24 total), there appeared to be a slightly skewed sex distribution within litters.

## 4. Discussion

Maternal health during pregnancy plays a critical role in shaping the future health and disease risk for offspring. There are dimensions of modern day life that challenge the ability to best manage stress which can lead to poor dietary habits and amplify existing psychosocial stresssors. While significant progress has been made in the preclinical space to understand the impact of individual exposures on maternal health, few studies have integrated multiple stressors to explore their combined effects. To date, there are no established animal models of maternal stress that have integrated multiple sources of environmental stress prior to conception and during pregnancy. In this study, we show that the combination of a chronic high fat diet and 6-weeks of social instability stress result in metabolic dysfunction, increased stress vulnerability and aberrant maternal behaviour.

Here, we found that female mice which had continuous access to a HF diet from 6 weeks of age and throughout pregnancy and lactation, demonstrated rapid body weight gain independent of SIS status. This physiological change was interesting to observe against the background of social stress exposure, given that stress-induced social subordination can profoundly influence eating behaviours and energy utilisation (Carneiro-Nascimento et al., 2020; Natterson-Horowitz and Cho, 2021). This was a result we had also observed amongst our pilot HF+/SIS+ cohort which had been exposed to 13-weeks of SIS in parallel with the same HF diet formulation (**Suppl. Fig. 1**). Although we could not quantify weekly food intake due to group housing conditions, we noted significant fluctuations in weight gain during SIS exposure, supporting our hypothesis that the social stress may have influenced eating patterns and energy expenditure. In regard to other maternal HF diet models in mice, several groups have observed consistent trends in increased body weight gain and an increase in various fat depots leading up to breeding (Bellisario et al., 2015, 2014; Bordeleau et al., 2020). The broad consensus from these studies is that the body weight increase may stem from increased daily consumption of the HF diet, due to its higher caloric content. This explanation is also supported by the observation that body weight returned to normal levels after the HF diet was discontinued (Bellisario et al., 2015, 2014).

Maternal HF diet models have predominantly served in elucidating the mechanisms underlying maternal obesity and the programming of obesity risk in offspring. Previous work has revealed maternal HF diet intake has induced a host of clinical features of metabolic syndrome including, insulin and leptin resistance, hypertension, fatty pancreas disease and transmission of nonalacoholic fatty liver disease to offspring (for review see: (Li et al., 2011)). We expected to find some degree of metabolic dysfunction as a consequence of HF diet feeding in both HF+ groups given that their weight gain tracked very closely together. However, statistically, we only observed significantly elevated fasting blood glucose and an impaired glucose response in HF+/SIS- mice relative to HF- controls, despite HF+/SIS+ mice showing similar values. These findings provide evidence that all HF+ mice show initial signs of a metabolic shift towards metabolic syndrome by 8-weeks of HF diet exposure which is consistent with previous iterations of maternal HF diet models in mice (Fernandes et al., 2021; Li et al., 2011; Moazzam et al., 2021; Murabayashi et al., 2013).

In the context of SIS in isolation, previous literature suggests there to be a consistent impact on HPA-axis function. This is largely characterised by elevated levels of corticosterone either during or at the end of the paradigm; or an increased corticosterone/ACTH ratio at the end of the paradigm (Koert et al., 2021b; Schmidt et al., 2010a; Yohn et al., 2019). The basis of the SIS model began in male mice which were found to be particularly sensitive to both the short-term and long-term effects of SIS on HPA-axis regulation (Chatterjee et al., 2009; Scharf et al., 2013; Schmidt et al., 2010b, 2007; Sterlemann et al., 2008). However, recent use of this paradigm in female mice has led to further insights into their vulnerability to chronic social stress-induced phenotypes. For example, one study that employed a 7-week SIS protocol in female mice, observed increased adrenal weight, decreased thymus weight, increased basal morning corticosterone and increased corticosterone/ACTH ratio at the end of the stress protocol (Schmidt et al., 2010a). In our stress paradigm, the adverse effects of SIS on the HPA-axis system were present exclusively in our HF+/SIS+ mice and absent in HF-/SIS+ mice. This highlights the significant role of macronutrient status in modulating the stress response, emphasising that a high-fat diet in this context primarily impedes stress adaptation. The added benefit of using the SIS paradigm in females is the ability to examine transgenerational transmission of these phenotypes to offspring (Saavedra-Rodríguez and Feig, 2013; Schmidt et al., 2010a; Yohn et al., 2019). In this regard, one study reported enhanced anxiety and social deficits transmitted to all F1 offspring coupled with elevated serum corticosterone levels (Saavedra-Rodríguez and Feig, 2013). Future studies are necessary to investigate the long-term impacts of chronic social stress and HF diet intake on the maternal stress response and how these factors may influence prenatal development.

At the behavioural level, we observed that neither HF diet or SIS alone or in combination appeared to substantially influence anxiety-like behaviours prior to conception. Yet, HF+/SIS+ mice did demonstrate increased locomotor activity based on the increase in total distance travelled during elevated plus maze testing. We attribute this to the differences in the experimental setup between the plus maze and the open field more than HF diet or stress exposure. In this respect, Bellisario et al., (2015) characterised HF diet fed dams as having lower locomotor activity 3 days prior to giving birth (Bellisario et al., 2015). However, this measurement was performed on pregnant dams and, perhaps unsurprisingly due to the additional weight gained, pregnancy has been shown to reduce levels of locomotor activity in mice (Martin-Fairey et al., 2019). This behavioural change to limit unnecessary movement could also be interpreted as being a protective mechanism for ensuring a successful delivery for both the dams and pups. Studies that have used the SIS paradigm in mice generally observe either an increase or no change in anxiety- related beahviour. However, there appear to be mixed behavioural profiles due to variations in the length of the paradigm, how soon behavioural tesing was performed following the protocol, cage density, strain, age and sex of mice used (for review see: (Koert et al., 2021a)). Yohn and colleagues exposed female adult C57Bl/6 mice to a 7-week SIS protocol with a cage change frequency of every 3 days (Yohn et al., 2019). They observed SIS mice travelled less distance in the open arms of the elevated plus maze and in the centre zone of the open field test. In contrast, another group found female CD1 mice displayed a less anxious behavioural profile and unaltered plasma corticosterone and ACTH levels following their 4-week, daily-SIS regimen (Dadomo et al., 2018). Interestingly, Schmidt et al., (2010) reported an increased corticosterone/ACTH ratio following their 7-week SIS paradigm in female CD1 mice, while locomotor activity during open field testing and time spent in the open arms of the elevated plus maze remained unchanged compared to controls (Schmidt et al., 2010a). Despite the lack of statistical significance in our plus maze data, the percentage of time spent in the open arms appears to be considerably less on average in the HF-/SIS+ group compared to other stress groups. We speculate that this measure did not reach statistical significance due to the behavioural variation within HF-/SIS+ and HF+/SIS+ mice. This variation could be explained by differing propensities for exploratory drive from a subset of mice inherently more susceptible to the influence of a long-term HF diet or SIS. While outside the scope of this study, the differing susceptibility to stress exposure is an area of active investigation and has been reported by others (Gururajan et al., 2019; Mueller et al., 2021; Pfau and Russo, 2015; Schmidt et al., 2010b).

Our last behavioural test to contextualise behaviour prior to inducing pregnancy was the nest building assay. Nest building is an inherent behaviour in rodents that is sensitive to changes in physiology and welfare (Neely et al., 2019). It has also been described that reproductive females are highly motivated to build high quality nests (Latham and Mason, 2004). Here, we observed nesting behaviours did not reach statistical significance as per nest scores across groups. Yet, we did observe a trend amongst both HF+ groups which demonstrated poorer nest quality on average compared to HF- groups. Further, our data also suggests that HF diet imparts a more pronounced impact on nesting behaviour as opposed to SIS. Whilst we acknowledge that performing this assay prior to, and after each stress intervention would have revealed more significant differences, our current observations are in line with Moazzam et al., (2021) that show pregnant HF diet mice had a reduced mean nest score suggesting that their nest building capacity was impaired (Moazzam et al., 2021). To our knowledge, nesting behaviours have not been assessed within the context of SIS paradigms, thus making our study the first to do so, albeit in the presence of a high-fat dietary regimen.

To further affirm our behavioural findings, we calculated an integrated anxiety- Z-score from our open field, elevated plus maze and nest building output (Guilloux et al., 2011; Kraeuter, 2023). As expected, the Z-score result reflected the behavioural variation we observed amongst mice within each stress group and did not show a consistent trend in either direction for an increase (> 0) or decrease (< 0) in anxiety across all 3 behavioural tests. This summary statistic reinforces that anxiety-like behaviours were not found to be impacted following the HF diet or SIS. To expand upon this behavioural profile, opting for more sensitive behavioural tests such as, the light-dark box test or tail suspension test may reveal more pronounced differences in anxiety-like behaviours.

Perhaps the most profound behavioural change we observed amongst these maternal stress groups occurred post-partum. We observed frequent episodes of maternal neglect, defined here as pup cannibalism shortly after birth, in both HF+ stress groups. We hypothesize that this aberrant maternal behavior could have an association with the aforementioned trend of poor nest quality amongst both HF+ groups. These results are consistent with those reported by Bellisario and colleagues (Bellisario et al., 2015, 2014) where they also found maternal HF diet feeding led to increased levels of cannibalistic behaviour. Due to such high levels of cannibalism following continuous feeding, they implemented a switch to a standard diet three days prior to delivery to promote pup survival. They also report HF diet dams displayed a greater latency time to approach their pups and more time spent self-grooming (Bellisario et al., 2015). Interestingly, our 6-week SIS protocol prior to pregnancy did not seem to produce a large effect on maternal care in the absence of a HF diet. Instead, we observed the highest frequency of care among HF-/SIS+ mice, whereas HF-/SIS- controls showed an even split in this aspect. However, we cannot entirely dismiss the influence of SIS on maternal care, considering that the LF diet includes a relatively high sucrose content, which might impact various aspects of maternal behavior that we did not measure here.

Lastly, the offspring phenotypical changes we report from these dams include but are not limited to, a marginal increase in body weight between HF+ and HF- offspring at the time of weaning (**Suppl. Table 1**). Many studies have shown in utero exposure to maternal HF diet or obesity to lead to adverse health outcomes in offspring; particularly, hyperphagia, increased adiposity and insulin resistance (Samuelsson et al., 2008; Wankhade et al., 2016). Moreover, we observed a skewed sex ratio favouring males within litters from HF+/SIS- dams, while HF+/SIS+ dams seemed to produce more females. There is limited literature in this area to explain this phenomenon but, observations from Rosenfeld et al., (2003) report similar findings using comparable dietary compositions. They assigned female mice to either a very high in saturated fat (VHF) or low saturated fat but high in carbohydrate (LF) diet throughout pregnancy and observed a large proportion of males born to VHF dams and much lower from LF dams (Rosenfeld et al., 2003). This is an area that should be explored further to better understand the influence of macronutrient intake during pregnancy on sex bias in offspring.

It is important to note that the primary technical limitation of this model is frequent cannibalistic behaviour due to continuous HF diet feeding, complicating the study of long-term impacts on prenatal development. Yet, modifying fat and sugar content during pregnancy and cross-fostering offspring represent acceptable methods to address this issue. Other limitations of our study include the exclusion of a standard chow diet control group. Our current HF- control group was fed a semi-purified diet, which matched the HF diet in sucrose and casein content, but contained a reduced amount of lard. Therefore, we cannot disregard the potential impact of high sucrose on maternal health and behaviour in this study. In addition, a more thorough metabolic assessment would have provided further insights into the role of metabolic dysfunction in these mice. Lastly, the addition of postpartum behavioural testing would have provided more functional context for how this combined stress impacts negative valence behaviours longer-term.

In conclusion, this is the first study to incorporate multiple sources of environmental stress into one female mouse model. This ‘two-hit’, HF+/SIS+ paradigm provides evidence that a maternal HF diet, in addition to social stress, seriously constrains stress adaptation. As a result of this failure to adapt to stress, these dams display negative postpartum maternal behaviour. This aberrant maternal behavior indicates a synergistic relationship between persistent overnourishment and chronic, unpredictable social stress, leading to prolonged dysregulation of the stress response. Future studies are warranted to further disentangle the relative impact of multiple stressors on maternal health, prenatal and postnatal development as well as investigate intervention strategies to mitigate negative outcomes.

## Supporting information

Supplemental statistical report 1-6

## Declaration of Generative AI and AI-assisted technologies in the writing process

During the preparation of this work the author(s) used ChatGPT in order to improve readability and refine sentence structure. After using this tool/service, the author(s) reviewed and edited the content as needed and take(s) full responsibility for the content of the publication.

## Data availability

Data will be made available upon request.

## Acknowledgements

This work was supported by a National Health and Medical Research Council (NHMRC) of Australia Investigator grant (APP1193648). The authors would like to acknowledge all animal monitoring and technical assistance provided from all Laboratory Animal Services (LAS) staff during this project. They also thank Dr. Stela Petkova for technical assistance with behavioural testing and optimisation. Fig 1A was created under a paid subscription for BioRender.com.

## Appendix A. Supplementary data

**Supplemental Figure 1.** Pilot data showing maternal HF diet induces rapid weight gain after 15-weeks of diet intake and 13-weeks of SIS.

**Supplemental Figure 2.** Pilot data showing no changes in anxiety-like behaviour via open field but only in elevated plus maze testing prior to pregnancy.

**Supplementary Table 1.** Reproductive outcomes summary.

**SupplementaryTable 2.** Chi-square frequency table of dams cannibalistic behaviour and sex distribution of live offspring within litters (Fig. 5 A-B).

**Supplemental Statistical Report 1** Fisher’s LSD post-hoc test following RM 2-way ANOVA of absolute weight gain over weeks 1-8 of diet intake (Fig 1B).

**Supplemental Statistical Report 2** Dunn’s post-hoc test followed by Friedman test for percentage of weight gained over weeks 1-8 of diet intake (Fig. 1C).

**Supplemental Statistical Report 3** Dunn’s post-hoc test followed by Kruskal-Wallis test for ipGTT response curve across stress groups (AUC) (Fig. 2B).

**Supplemental Statistical Report 4** Dunn’s post-hoc test followed by Kruskal-Wallis test for fasting blood glucose prior to ipGTT (Fig. 2C).

**Supplemental Statistical Report 5** Fisher’s LSD post-hoc test followed by 2way RM ANOVA for plasma adrenocorticotropic hormone at baseline and post SIS (Fig. 3A).

**Supplemental Statistical Report 6** Tukey’s post-hoc test followed by 2way RM ANOVA for plasma corticosterone across stress groups between baseline and post SIS (Fig. 3B).

## Author contributions

Morgan C Bucknor conceived experimental design, conducted experiments, performed data analysis and wrote the original draft of the manuscript. Anand Gururajan assisted with experimental design, reviewed and edited subsequent drafts of the manuscript. Russell C Dale acquired funding, assisted with data interpretation and edited subsequent drafts of the manuscript. Markus J Hofer conceived and supervised experiments, reviewed and edited subsequent drafts of the manuscript.

## References

Bartolomucci, A., Palanza, P., Sacerdote, P., Panerai, A.E., Sgoifo, A., Dantzer, R., Parmigiani, S., 2005. Social factors and individual vulnerability to chronic stress exposure. Neurosci. Biobehav. Rev. 29, 67–81. 10.1016/j.neubiorev.2004.06.009

Bellisario, V., Berry, A., Capoccia, S., Raggi, C., Panetta, P., Branchi, I., Piccaro, G., Giorgio, M., Pelicci, P.G., Cirulli, F., 2014. Gender-dependent resiliency to stressful and metabolic challenges following prenatal exposure to high-fat diet in the p66Shc-/- mouse. Front. Behav. Neurosci. 8, 1–12. 10.3389/fnbeh.2014.00285

Bellisario, V., Panetta, P., Balsevich, G., Baumann, V., Noble, J., Raggi, C., Nathan, O., Berry, A., Seckl, J., Schmidt, M., Holmes, M., Cirulli, F., 2015. Maternal high-fat diet acts as a stressor increasing maternal glucocorticoids’ signaling to the fetus and disrupting maternal behavior and brain activation in C57BL/6J mice. Psychoneuroendocrinology 60, 138–150. 10.1016/j.psyneuen.2015.06.012

Bordeleau, M., Lacabanne, C., Cossío, L.F. de, Vernoux, N., Savage, J., González- Ibáñez, F., Tremblay, M.-E., 2020. Microglial and peripheral immune priming is partially sexually dimorphic in adolescent mouse offspring exposed to maternal high-fat diet. J. Neuroinflammation 17, 1–28. 10.21203/rs.3.rs-25000/v1

Bronson, S.L., Bale, T.L., 2016. The Placenta as a Mediator of Stress Effects on Neurodevelopmental Reprogramming. Neuropsychopharmacology 41, 207–218. 10.1038/npp.2015.231

Carneiro-Nascimento, S., Opacka-Juffry, J., Costabile, A., Boyle, C.N., Herde, A.M., Ametamey, S.M., Sigrist, H., Pryce, C.R., Patterson, M., 2020. Chronic social stress in mice alters energy status including higher glucose need but lower brain utilization. Psychoneuroendocrinology 119. 10.1016/j.psyneuen.2020.104747

Chatterjee, O., Taylor, L.A., Ahmed, S., Nagaraj, S., Hall, J.J., Finckbeiner, S.M., Chan, P.S., Suda, N., King, J.T., Zeeman, M.L., McCobb, D.P., 2009. Social stress alters expression of large conductance calcium-activated potassium channel subunits in mouse adrenal medulla and pituitary glands. J. Neuroendocrinol. 21, 167–176. 10.1111/j.1365-2826.2009.01823.x

Cirulli, F., Musillo, C., Berry, A., 2020. Maternal Obesity as a Risk Factor for Brain Development and Mental Health in the Offspring. Neuroscience 447, 122–135. 10.1016/j.neuroscience.2020.01.023

Dadomo, H., Gioiosa, L., Cigalotti, J., Ceresini, G., Parmigiani, S., Palanza, P., 2018. What is stressful for females? Differential effects of unpredictable environmental or social stress in CD1 female mice. Horm. Behav. 98, 22–32. 10.1016/j.yhbeh.2017.11.013

Deacon, R.M.J., 2006. Assessing nest building in mice. Nat. Protoc. 1, 1117–1119. 10.1038/nprot.2006.170

Driscoll, A.K., Gregory, E.C.W., 2020. Increases in Prepregnancy Obesity: United States, 2016-2019. NCHS Data Brief 1–8.

Fernandes, D.J., Spring, S., Roy, A.R., Qiu, L.R., Yee, Y., Nieman, B.J., Lerch, J.P., Palmert, M.R., 2021. Exposure to maternal high-fat diet induces extensive changes in the brain of adult offspring. Transl. Psychiatry 11, 1–9. 10.1038/s41398-021-01274-1

Guilloux, J.P., Seney, M., Edgar, N., Sibille, E., 2011. Integrated behavioral z-scoring increases the sensitivity and reliability of behavioral phenotyping in mice: Relevance to emotionality and sex. J. Neurosci. Methods 197, 21–31. 10.1016/j.jneumeth.2011.01.019

Gururajan, A., van de Wouw, M., Boehme, M., Becker, T., O’Connor, R., Bastiaanssen, T.F.S., Moloney, G.M., Lyte, J.M., Ventura Silva, A.P., Merckx, B., Dinan, T.G., Cryan, J.F., 2019. Resilience to chronic stress is associated with specific neurobiological, neuroendocrine and immune responses. Brain. Behav. Immun. 80, 583–594. 10.1016/j.bbi.2019.05.004

Hall, K.D., 2018. Did the Food Environment Cause the Obesity Epidemic? Obesity 26, 11–13. 10.1002/oby.22073

Han, V.X., Patel, S., Jones, H.F., Dale, R.C., 2021a. Maternal immune activation and neuroinflammation in human neurodevelopmental disorders. Nat. Rev. Neurol. 17, 564–579. 10.1038/s41582-021-00530-8

Han, V.X., Patel, S., Jones, H.F., Dale, R.C., 2021b. Maternal immune activation and neuroinflammation in human neurodevelopmental disorders. Nat. Rev. Neurol. 0123456789. 10.1038/s41582-021-00530-8

Herman, J.P., 2013. Neural control of chronic stress adaptation. Front. Behav. Neurosci. 7, 1–12. 10.3389/fnbeh.2013.00061

Hu, S., Wang, L., Yang, D., Li, L., Togo, J., Wu, Y., Liu, Q., Li, B., Li, M., Wang, G., Zhang, X., Niu, C., Li, J., Xu, Y., Couper, E., Whittington-Davies, A., Mazidi, M., Luo, L., Wang, S., Douglas, A., Speakman, J.R., 2018. Dietary Fat, but Not Protein or Carbohydrate, Regulates Energy Intake and Causes Adiposity in Mice. Cell Metab. 28, 415–431.e4. 10.1016/j.cmet.2018.06.010

Koert, A., Ploeger, A., Bockting, C.L.H., Schmidt, M. V., Lucassen, P.J., Schrantee, A., Mul, J.D., 2021a. The social instability stress paradigm in rat and mouse: A systematic review of protocols, limitations, and recommendations. Neurobiol. Stress 15, 100410. 10.1016/j.ynstr.2021.100410

Koert, A., Ploeger, A., Bockting, C.L.H., Schmidt, M. V., Lucassen, P.J., Schrantee, A., Mul, J.D., 2021b. The social instability stress paradigm in rat and mouse: A systematic review of protocols, limitations, and recommendations. Neurobiol. Stress 15, 100410. 10.1016/j.ynstr.2021.100410

Kraeuter, A.K., 2023. The use of integrated behavioural z-scoring in behavioural neuroscience – A perspective article. J. Neurosci. Methods 384, 109751. 10.1016/j.jneumeth.2022.109751

Latham, N., Mason, G., 2004. From house mouse to mouse house: The behavioural biology of free-living Mus musculus and its implications in the laboratory. Appl. Anim. Behav. Sci. 86, 261–289. 10.1016/j.applanim.2004.02.006

Li, M., Sloboda, D.M., Vickers, M.H., 2011. Maternal obesity and developmental programming of metabolic disorders in offspring: Evidence from animal models. Exp. Diabetes Res. 2011. 10.1155/2011/592408

Martin-Fairey, C.A., Zhao, P., Wan, L., Roenneberg, T., Fay, J., Ma, X., McCarthy, R., Jungheim, E.S., England, S.K., Herzog, E.D., 2019. Pregnancy Induces an Earlier Chronotype in Both Mice and Women. J. Biol. Rhythms 34, 323–331. 10.1177/0748730419844650

McEwen, B.S., Wingfield, J.C., 2003. The concept of allostasis in biology and biomedicine. Horm. Behav. 43, 2–15. 10.1016/S0018-506X(02)00024-7

Moazzam, S., Jarmasz, J.S., Jin, Y., Siddiqui, T.J., Cattini, P.A., 2021. Effects of high fat diet-induced obesity and pregnancy on prepartum and postpartum maternal mouse behavior. Psychoneuroendocrinology 126, 105147. 10.1016/j.psyneuen.2021.105147

Mueller, F.S., Scarborough, J., Schalbetter, S.M., Richetto, J., Kim, E., Couch, A., Yee, Y., Lerch, J.P., Vernon, A.C., Weber-Stadlbauer, U., Meyer, U., 2021. Behavioral, neuroanatomical, and molecular correlates of resilience and susceptibility to maternal immune activation. Mol. Psychiatry 26, 396–410. 10.1038/s41380-020-00952-8

Mumtaz, F., Khan, M.I., Zubair, M., Dehpour, A.R., 2018. Neurobiology and consequences of social isolation stress in animal model—A comprehensive review. Biomed. Pharmacother. 105, 1205–1222. 10.1016/j.biopha.2018.05.086

Murabayashi, N., Sugiyama, T., Zhang, L., Kamimoto, Y., Umekawa, T., Ma, N., Sagawa, N., 2013. Maternal high-fat diets cause insulin resistance through inflammatory changes in fetal adipose tissue. Eur. J. Obstet. Gynecol. Reprod. Biol. 169, 39–44. 10.1016/j.ejogrb.2013.02.003

Natterson-Horowitz, B., Cho, J.H., 2021. Stress, Subordination, and Anomalies of Feeding Across the Tree of Life: Implications for Interpreting Human Eating Disorders. Front. Psychol. 12. 10.3389/fpsyg.2021.727554

Neely, C.L.C., Pedemonte, K.A., Boggs, K.N., Flinn, J.M., 2019. Nest building behavior as an early indicator of behavioral deficits in mice. J. Vis. Exp. 2019, 1–8. 10.3791/60139

Pfau, M.L., Russo, S.J., 2015. Peripheral and central mechanisms of stress resilience. Neurobiol. Stress 1, 66–79. 10.1016/j.ynstr.2014.09.004

Radley, J.J., Herman, J.P., 2023. Preclinical Models of Chronic Stress: Adaptation or Pathology? Biol. Psychiatry 94, 194–202. 10.1016/j.biopsych.2022.11.004

Ramsay, D.S., Woods, S.C., 2014. Clarifying the roles of homeostasis and allostasis in physiological regulation. Psychol. Rev. 121, 225–247. 10.1037/a0035942

Rosenfeld, C.S., Grimm, K.M., Livingston, K.A., Brokman, A.M., Lamberson, W.E., Roberts, R.M., 2003. Striking variation in the sex ratio of pups born to mice according to whether maternal diet is high in fat or carbohydrate. Proc. Natl. Acad. Sci. U. S. A. 100, 4628–4632. 10.1073/pnas.0330808100

Saavedra-Rodríguez, L., Feig, L.A., 2013. Chronic social instability induces anxiety and defective social interactions across generations. Biol. Psychiatry 73, 44–53. 10.1016/j.biopsych.2012.06.035

Samuelsson, A.M., Matthews, P.A., Argenton, M., Christie, M.R., McConnell, J.M., Jansen, E.H.J.M., Piersma, A.H., Ozanne, S.E., Twinn, D.F., Remacle, C., Rowlerson, A., Poston, L., Taylor, P.D., 2008. Diet-induced obesity in female mice leads to offspring hyperphagia, adiposity, hypertension, and insulin resistance: A novel murine model of developmental programming. Hypertension 51, 383–392. 10.1161/HYPERTENSIONAHA.107.101477

Scharf, S.H., Sterlemann, V., Liebl, C., Müller, M.B., Schmidt, M. V., 2013. Chronic social stress during adolescence: Interplay of paroxetine treatment and ageing. Neuropharmacology 72, 38–46. 10.1016/j.neuropharm.2013.03.035

Schmidt, M. V., Scharf, S.H., Liebl, C., Harbich, D., Mayer, B., Holsboer, F., Müller, M.B., 2010a. A novel chronic social stress paradigm in female mice. Horm. Behav. 57, 415–420. 10.1016/j.yhbeh.2010.01.010

Schmidt, M. V., Scharf, S.H., Sterlemann, V., Ganea, K., Liebl, C., Holsboer, F., Müller, M.B., 2010b. High susceptibility to chronic social stress is associated with a depression-like phenotype. Psychoneuroendocrinology 35, 635–643. 10.1016/j.psyneuen.2009.10.002

Schmidt, M. V., Sterlemann, V., Ganea, K., Liebl, C., Alam, S., Harbich, D., Greetfeld, M., Uhr, M., Holsboer, F., Müller, M.B., 2007. Persistent neuroendocrine and behavioral effects of a novel, etiologically relevant mouse paradigm for chronic social stress during adolescence. Psychoneuroendocrinology 32, 417–429. 10.1016/j.psyneuen.2007.02.011

Sterlemann, V., Ganea, K., Liebl, C., Harbich, D., Alam, S., Holsboer, F., Müller, M.B., Schmidt, M. V., 2008. Long-term behavioral and neuroendocrine alterations following chronic social stress in mice: Implications for stress-related disorders. Horm. Behav. 53, 386–394. 10.1016/j.yhbeh.2007.11.001

Wankhade, U.D., Thakali, K.M., Shankar, K., 2016. Persistent influence of maternal obesity on offspring health: Mechanisms from animal models and clinical studies. Mol. Cell. Endocrinol. 435, 7–19. 10.1016/j.mce.2016.07.001

Yohn, C.N., Ashamalla, S.A., Bokka, L., Gergues, M.M., Garino, A., Samuels, B.A., 2019. Social instability is an effective chronic stress paradigm for both male and female mice. Neuropharmacology 160, 107780. 10.1016/j.neuropharm.2019.107780

Ziauddeen, N., Huang, J.Y., Taylor, E., Roderick, P.J., Godfrey, K.M., Alwan, N.A., 2022. Interpregnancy weight gain and childhood obesity: analysis of a UK population-based cohort. Int. J. Obes. 46, 211–219. 10.1038/s41366-021-00979-z

