## Supplemental Figure 1 for "High fat diet consumption and social instability stress impair stress adaptation and maternal care in C57Bl/6 mice"

Supplementary Material 2 – Figures

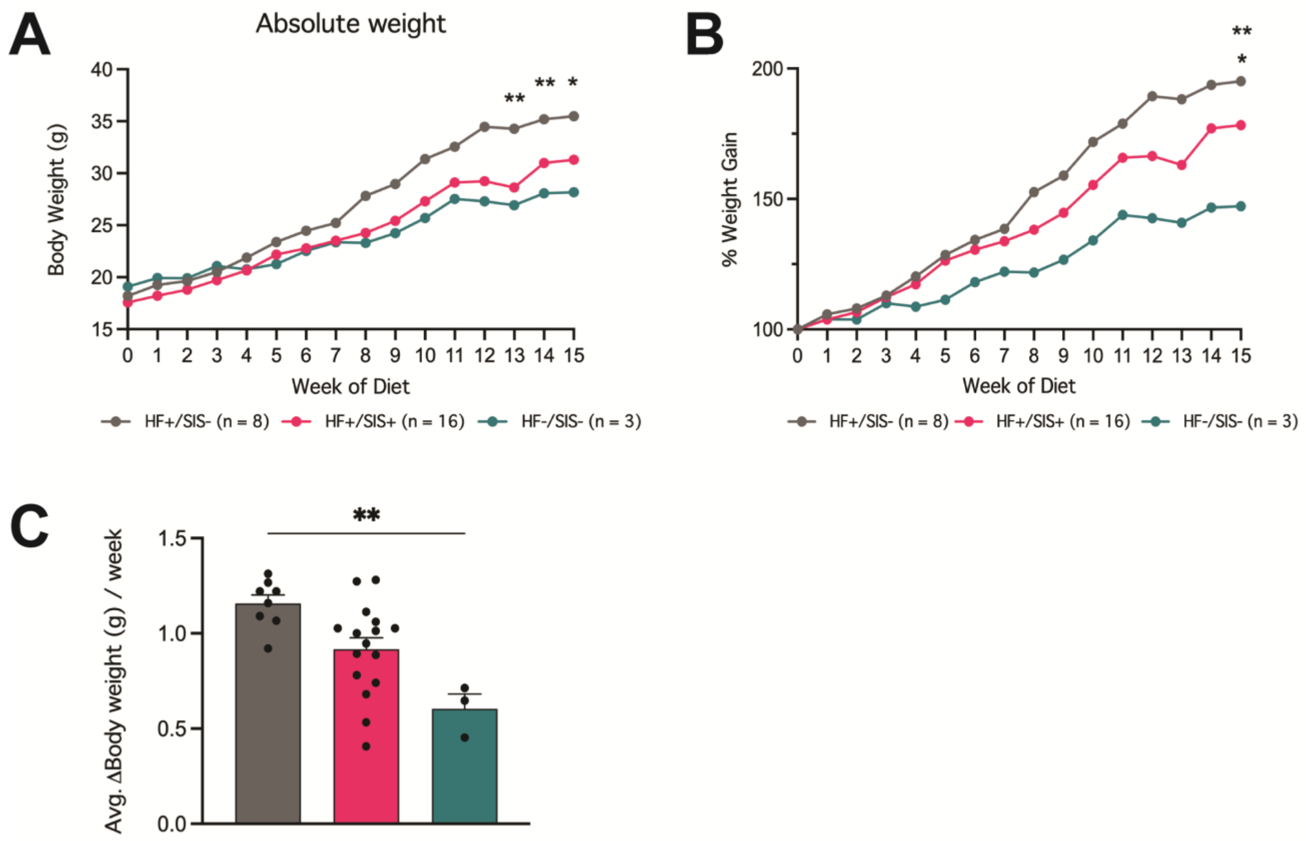


**Supplemental Figure 1. Pilot data showing maternal HF diet induces rapid weight gain after 15-weeks of diet intake and 13-weeks of SIS.** (A) Absolute weight gain across 15 weeks of respective dietary intake. (B) Percentage of weight gained over 15 weeks of feeding in parallel with SIS. (C) Average change in body weight per week during HF diet feeding and/or SIS. Data shown as mean ± SEM. *Denotes statistically significant difference (*p<0.05, **p<0.01). Fig A-B: 2Way RM ANOVA; Tukey's post hoc test. Fig C: Kruskal Wallis; Dunn's post hoc test.

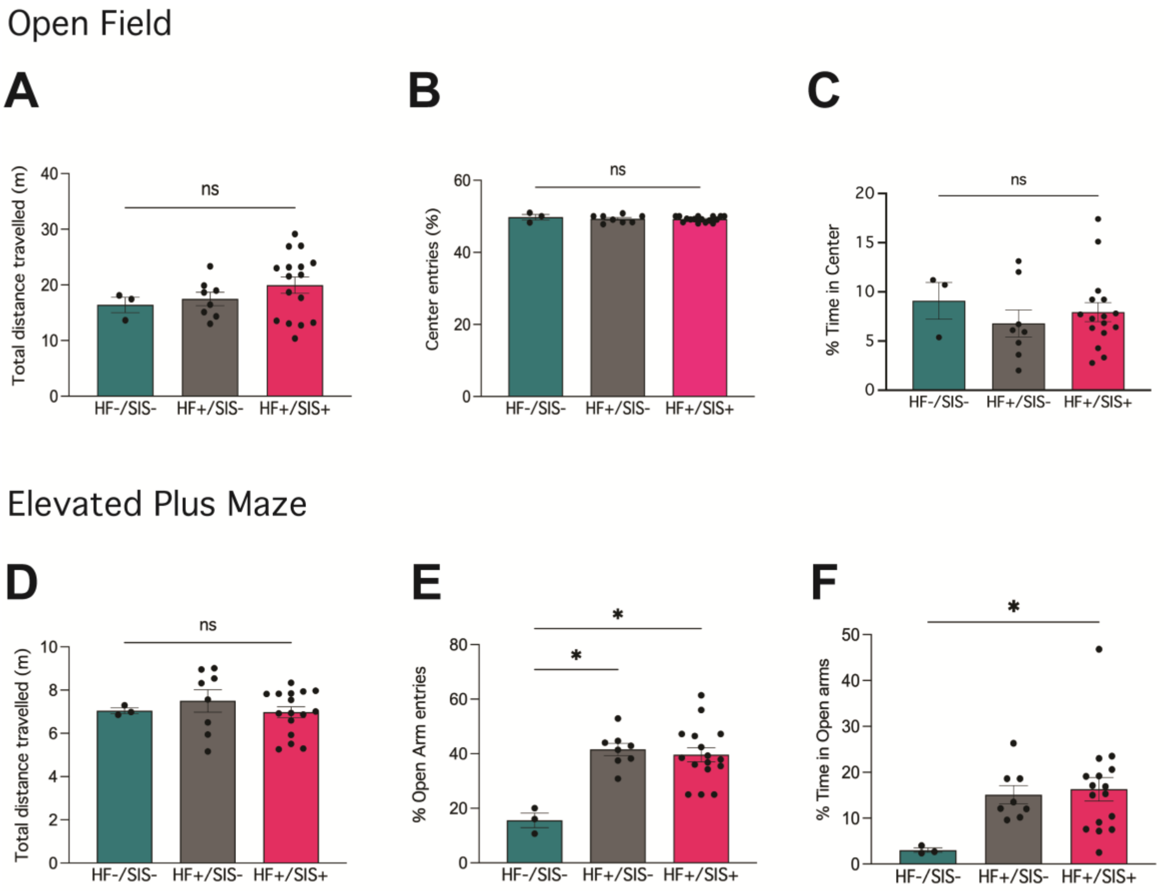


**Supplemental Figure 2. Pilot data showing no changes in anxiety-like behaviour via open field but only in elevated plus maze testing prior to pregnancy.** (A) Total distance travelled in open field arena (B) Percentage of entries into center zone (C) Percentage of time in center zone (D) Total distance travelled during elevated plus maze test (E) Percentage of entries into the open arms (F) Percentage of time spent in the open arms. Data shown as mean ± SEM. *Denotes statistically significant difference: (*p<0.05). Fig A-F: Kruskall wallis; Dunn's post hoc test.
