## Supplemental statistical report 1-6 for "High fat diet consumption and social instability stress impair stress adaptation and maternal care in C57Bl/6 mice"

Supplementary Material 1 – Tables and Statistical Reports

**Supplementary Table 1.** Reproductive outcomes summary.

| Reproductive Outcome | Measurement | Maternal stress group | | | |
|---|---|---|---|---|---|
| | | HF-/SIS- | HF-/SIS+ | HF+/SIS- | HF+/SIS+ |
| Litter size | Pups born (n) | 7.5 ± 0.28 | 5.4 ± 0.68 | 6.5 ± 1.5 | 5.3 ± 0.33 |
| Sex ratio | Male pups (n) | 3.25 ± 0.25 | 3 ± 0.44 | 5 ± 3 | 2 ± 0.57 |
| | Female pups (n) | 4.25 ± 0.48 | 2.4 ± 0.6 | 1.5 ± 1.5 | 3.3 ± 0.33 |
| Weanling weight (PnD 28-30) | Male weight (g) | 12.7 ± 0.33 | 14.5 ± 0.32** | 14.1 ± 0.2* | 14.25 ± 0.81 |
| | Female weight (g) | 11.8 ± 0.25 | 13.3 ± 0.28** | 13 ± 0.3 | 14.35 ± 0.36*** |

Data shown as mean ± SEM.

(*p<0.05), (**p<0.01), (***p<0.001), relative to HF-/SIS- control group following ANOVA and Dunnett's post-hoc test.

PnD = postnatal day.

**Supplementary Table 2.** Chi-square frequency table of dams' cannibalistic behaviour and sex distribution of live offspring within litters.

| Maternal Behaviour | HF-/SIS- (n = 8) | HF-/SIS+ (n = 14) | HF+/SIS- (n = 7) | HF+/SIS+ (n = 16) | p-Value |
|---|---|---|---|---|---|
| Cannibalism | 50% | 29% | 71% | 81% | <0.0001[a] |
| No cannibalism | 50% | 71% | 29% | 19% | |
| **Sex of live pups** | **HF-/SIS- (n = 30)** | **HF-/SIS+ (n = 54)** | **HF+/SIS- (n = 13)** | **HF+/SIS+ (n = 16)** | <0.0001[b] |
| Male | 43% | 56% | 77% | 38% | |
| Female | 57% | 44% | 23% | 62% | |

Data shown as absolute proportions.

[a]Chi-square test was applied to compare the frequency of maternal care versus neglect (cannibalism).

[b]Chi-square test was applied to compare the frequency of male versus female live offspring.

**Supplemental Statistical Report 1.** Fisher's LSD post-hoc test following RM 2-way ANOVA of absolute weight gain over weeks 1-8 of diet intake (Fig 1B).

| Comparison | Mean 1 | Mean 2 | Mean diff. | SE of diff. | Summary | Individual P-value |
|---|---|---|---|---|---|---|
| **Week 1** | | | | | | |
| HF+ / SIS+ vs. HF+ / SIS- | 17.56 | 17.80 | -0.2375 | 0.4991 | ns | 0.6407 |
| HF+ / SIS+ vs. HF- / SIS- | 17.56 | 16.50 | 1.063 | 0.4688 | * | 0.0364 |
| HF+ / SIS+ vs. HF- / SIS+ | 17.56 | 17.01 | 0.5500 | 0.3783 | ns | 0.1578 |
| HF+ / SIS- vs. HF- / SIS- | 17.80 | 16.50 | 1.300 | 0.5230 | * | 0.0264 |
| HF+ / SIS- vs. HF- / SIS+ | 17.80 | 17.01 | 0.7875 | 0.4438 | ns | 0.1027 |
| HF- / SIS- vs. HF- / SIS+ | 16.50 | 17.01 | -0.5125 | 0.4094 | ns | 0.2338 |
| **Week 2** | | | | | | |
| HF+ / SIS+ vs. HF+ / SIS- | 17.85 | 18.60 | -0.7500 | 0.5958 | ns | 0.2319 |
| HF+ / SIS+ vs. HF- / SIS- | 17.85 | 16.79 | 1.063 | 0.4627 | * | 0.0346 |
| HF+ / SIS+ vs. HF- / SIS+ | 17.85 | 17.56 | 0.2875 | 0.3959 | ns | 0.4735 |
| HF+ / SIS- vs. HF- / SIS- | 18.60 | 16.79 | 1.813 | 0.6203 | * | 0.0125 |
| HF+ / SIS- vs. HF- / SIS+ | 18.60 | 17.56 | 1.038 | 0.5722 | ns | 0.0983 |
| HF- / SIS- vs. HF- / SIS+ | 16.79 | 17.56 | -0.7750 | 0.4319 | ns | 0.0936 |
| **Week 3** | | | | | | |
| HF+ / SIS+ vs. HF+ / SIS- | 19.64 | 20.34 | -0.7000 | 0.5719 | ns | 0.2470 |
| HF+ / SIS+ vs. HF- / SIS- | 19.64 | 17.54 | 2.100 | 0.3986 | **** | <0.0001 |
| HF+ / SIS+ vs. HF- / SIS+ | 19.64 | 18.56 | 1.075 | 0.3609 | ** | 0.0057 |
| HF+ / SIS- vs. HF- / SIS- | 20.34 | 17.54 | 2.800 | 0.5919 | *** | 0.0006 |
| HF+ / SIS- vs. HF- / SIS+ | 20.34 | 18.56 | 1.775 | 0.5672 | * | 0.0101 |
| HF- / SIS- vs. HF- / SIS+ | 17.54 | 18.56 | -1.025 | 0.3917 | * | 0.0185 |
| **Week 4** | | | | | | |
| HF+/SIS+ vs. HF+/SIS- | 20.63 | 21.36 | -0.7312 | 0.4894 | ns | 0.1580 |
| HF+/SIS+ vs. HF-/SIS- | 20.63 | 18.31 | 2.319 | 0.3275 | **** | <0.0001 |
| HF+/SIS+ vs. HF-/SIS+ | 20.63 | 19.24 | 1.394 | 0.3898 | ** | 0.0012 |
| HF+/SIS- vs. HF-/SIS- | 21.36 | 18.31 | 3.050 | 0.4421 | **** | <0.0001 |
| HF+/SIS- vs. HF-/SIS+ | 21.36 | 19.24 | 2.125 | 0.4901 | *** | 0.0007 |
| HF-/SIS- vs. HF-/SIS+ | 18.31 | 19.24 | -0.9250 | 0.3285 | * | 0.0101 |
| **Week 5** | | | | | | |
| HF+ / SIS+ vs. HF+ / SIS- | 21.38 | 21.91 | -0.5375 | 0.7162 | ns | 0.4683 |
| HF+ / SIS+ vs. HF- / SIS- | 21.38 | 18.83 | 2.550 | 0.4195 | **** | <0.0001 |
| HF+ / SIS+ vs. HF- / SIS+ | 21.38 | 19.62 | 1.756 | 0.4373 | *** | 0.0004 |
| HF+ / SIS- vs. HF- / SIS- | 21.91 | 18.83 | 3.088 | 0.6727 | ** | 0.0013 |
| HF+ / SIS- vs. HF- / SIS+ | 21.91 | 19.62 | 2.294 | 0.6840 | ** | 0.0077 |
| HF- / SIS- vs. HF- / SIS+ | 18.83 | 19.62 | -0.7937 | 0.3616 | * | 0.0398 |
| **Week 6** | | | | | | |
| HF+ / SIS+ vs. HF+ / SIS- | 21.89 | 22.90 | -1.006 | 0.9872 | ns | 0.3322 |
| HF+ / SIS+ vs. HF- / SIS- | 21.89 | 18.94 | 2.956 | 0.4795 | **** | <0.0001 |
| HF+ / SIS+ vs. HF- / SIS+ | 21.89 | 19.91 | 1.981 | 0.4780 | *** | 0.0003 |
| HF+ / SIS- vs. HF- / SIS- | 22.90 | 18.94 | 3.963 | 0.9360 | ** | 0.0028 |
| HF+ / SIS- vs. HF- / SIS+ | 22.90 | 19.91 | 2.988 | 0.9352 | * | 0.0125 |
| HF- / SIS- vs. HF- / SIS+ | 18.94 | 19.91 | -0.9750 | 0.3605 | * | 0.0141 |
| **Week 7** | | | | | | |
| HF+ / SIS+ vs. HF+ / SIS- | 22.54 | 23.23 | -0.6875 | 0.9350 | ns | 0.4756 |
| HF+ / SIS+ vs. HF- / SIS- | 22.54 | 19.56 | 2.975 | 0.5876 | **** | <0.0001 |
| HF+ / SIS+ vs. HF- / SIS+ | 22.54 | 20.65 | 1.888 | 0.5434 | ** | 0.0023 |
| HF+ / SIS- vs. HF- / SIS- | 23.23 | 19.56 | 3.663 | 0.8533 | ** | 0.0019 |
| HF+ / SIS- vs. HF- / SIS+ | 23.23 | 20.65 | 2.575 | 0.8235 | * | 0.0138 |
| HF- / SIS- vs. HF- / SIS+ | 19.56 | 20.65 | -1.088 | 0.3864 | * | 0.0137 |
| **Week 8** | | | | | | |
| HF+ / SIS+ vs. HF+ / SIS- | 24.25 | 24.30 | -0.05000 | 1.273 | ns | 0.9693 |
| HF+ / SIS+ vs. HF- / SIS- | 24.25 | 19.49 | 4.763 | 0.7060 | **** | <0.0001 |

| | | | | | | |
|---|---|---|---|---|---|---|
| HF+ / SIS+ vs. HF- / SIS+ | 24.25 | 20.31 | 3.944 | 0.7285 | **** | <0.0001 |
| HF+ / SIS- vs. HF- / SIS- | 24.30 | 19.49 | 4.813 | 1.124 | ** | 0.0028 |
| HF+ / SIS- vs. HF- / SIS+ | 24.30 | 20.31 | 3.994 | 1.139 | ** | 0.0076 |
| HF- / SIS- vs. HF- / SIS+ | 19.49 | 20.31 | -0.8187 | 0.4188 | ns | 0.0639 |

**Supplemental Statistical Report 2.** Dunn's post-hoc test followed by Friedman test for percentage of weight gained over weeks 1-8 of diet intake (Fig. 1C).

| Comparison | Rank sum 1 | Rank sum 2 | Rank sum diff. | Summary | Adjusted P-value |
|---|---|---|---|---|---|
| HF+ / SIS+ vs. HF+ / SIS- | 27.50 | 33.50 | -6.000 | ns | >0.9999 |
| HF+ / SIS+ vs. HF- / SIS- | 27.50 | 14.50 | 13.00 | ns | 0.1057 |
| HF+ / SIS+ vs. HF- / SIS+ | 27.50 | 14.50 | 13.00 | ns | 0.1057 |
| HF+ / SIS- vs. HF- / SIS- | 33.50 | 14.50 | 19.00 | ** | 0.0031 |
| HF+ / SIS- vs. HF- / SIS+ | 33.50 | 14.50 | 19.00 | ** | 0.0031 |
| HF- / SIS- vs. HF- / SIS+ | 14.50 | 14.50 | 0.000 | ns | >0.9999 |
| HF+ / SIS+ vs. HF+ / SIS- | 27.50 | 33.50 | -6.000 | ns | >0.9999 |

**Supplemental Statistical Report 3.** Dunn's post-hoc test followed by Kruskal-Wallis test for ipGTT response curve across stress groups (AUC) (Fig. 2B).

| Comparison | Mean rank 1 | Mean rank 2 | Mean rank diff. | Summary | Adjusted P-value |
|---|---|---|---|---|---|
| HF+ / SIS+ vs. HF- / SIS+ | 20.56 | 9.813 | 10.75 | ns | 0.1314 |
| HF+ / SIS+ vs. HF+ / SIS- | 20.56 | 24.50 | -3.938 | ns | >0.9999 |
| HF+ / SIS+ vs. HF- / SIS- | 20.56 | 11.13 | 9.438 | ns | 0.2652 |
| HF- / SIS+ vs. HF+ / SIS- | 9.813 | 24.50 | -14.69 | * | 0.0104 |
| HF- / SIS+ vs. HF- / SIS- | 9.813 | 11.13 | -1.313 | ns | >0.9999 |
| HF+ / SIS- vs. HF- / SIS- | 24.50 | 11.13 | 13.38 | * | 0.0261 |
| HF+ / SIS+ vs. HF- / SIS+ | 20.56 | 9.813 | 10.75 | ns | 0.1314 |

**Supplemental Statistical Report 4.** Dunn's post-hoc test followed by Kruskal-Wallis test for fasting blood glucose prior to ipGTT (Fig. 2C).

| Comparison | Mean rank 1 | Mean rank 2 | Mean rank diff. | Summary | Adjusted P-value |
|---|---|---|---|---|---|
| HF+ / SIS+ vs. HF- / SIS+ | 20.50 | 12.25 | 8.250 | ns | 0.4704 |
| HF+ / SIS+ vs. HF+ / SIS- | 20.50 | 23.50 | -3.000 | ns | >0.9999 |
| HF+ / SIS+ vs. HF- / SIS- | 20.50 | 9.750 | 10.75 | ns | 0.1310 |
| HF- / SIS+ vs. HF+ / SIS- | 12.25 | 23.50 | -11.25 | ns | 0.0984 |
| HF- / SIS+ vs. HF- / SIS- | 12.25 | 9.750 | 2.500 | ns | >0.9999 |
| HF+ / SIS- vs. HF- / SIS- | 23.50 | 9.750 | 13.75 | * | 0.0201 |

| HF+ / SIS+ vs. HF- / SIS+ | 20.50 | 12.25 | 8.250 | ns | 0.4704 |
|---|---|---|---|---|---|

**Supplemental Statistical Report 5.** Fisher's LSD post-hoc test followed by 2way RM ANOVA for plasma adrenocorticotropic hormone at baseline and post SIS (Fig. 3A).

| Comparison | Predicted mean 1 | Predicted mean 2 | Predicted mean diff. | t | DF | Summary | Individual P-value |
|---|---|---|---|---|---|---|---|
| HF- / SIS- Baseline vs Post SIS | 193.4 | 83.86 | 109.5 | 2.191 | 36 | * | 0.0350 |
| HF- / SIS+ Baseline vs Post SIS | 222.6 | 101.3 | 121.3 | 2.972 | 36 | ** | 0.0053 |
| HF+ / SIS- Baseline vs Post SIS | 189.9 | 93.05 | 96.81 | 1.936 | 36 | ns | 0.0607 |
| HF+ / SIS+ Baseline vs Post SIS | 245.4 | 209.4 | 35.99 | 0.8817 | 36 | ns | 0.3838 |
| **Baseline** | | | | | | | |
| HF- / SIS- vs HF- / SIS+ | 193.4 | 222.6 | -29.24 | 0.5715 | 72 | ns | 0.5695 |
| HF- / SIS- vs HF+ / SIS- | 193.4 | 189.9 | 3.525 | 0.06289 | 72 | ns | 0.9500 |
| HF- / SIS- vs HF+ / SIS+ | 193.4 | 245.4 | -51.97 | 1.016 | 72 | ns | 0.3131 |
| HF- / SIS+ vs HF+ / SIS- | 222.6 | 189.9 | 32.77 | 0.6404 | 72 | ns | 0.5240 |
| HF- / SIS+ vs HF+ / SIS+ | 222.6 | 245.4 | -22.73 | 0.4967 | 72 | ns | 0.6209 |
| HF+ / SIS- vs HF+ / SIS+ | 189.9 | 245.4 | -55.50 | 1.085 | 72 | ns | 0.2817 |
| **Post SIS** | | | | | | | |
| HF- / SIS- vs HF- / SIS+ | 83.86 | 101.3 | -17.47 | 0.3414 | 72 | ns | 0.7338 |
| HF- / SIS- vs HF+ / SIS- | 83.86 | 93.05 | -9.195 | 0.1641 | 72 | ns | 0.8701 |
| HF- / SIS- vs HF+ / SIS+ | 83.86 | 209.4 | -125.5 | 2.453 | 72 | * | 0.0166 |
| HF- / SIS+ vs HF+ / SIS- | 101.3 | 93.05 | 8.272 | 0.1617 | 72 | ns | 0.8720 |
| HF- / SIS+ vs HF+ / SIS+ | 101.3 | 209.4 | -108.0 | 2.361 | 72 | * | 0.0209 |
| HF+ / SIS- vs HF+ / SIS+ | 93.05 | 209.4 | -116.3 | 2.273 | 72 | * | 0.0260 |

**Supplemental Statistical Report 6.** Tukey's post-hoc test followed by 2way RM ANOVA for plasma corticosterone across stress groups between baseline and post SIS (Fig. 3B).

| Comparison | Predicted mean 1 | Predicted mean 2 | Predicted mean diff. | q | DF | Summary | Adjusted P-value |
|---|---|---|---|---|---|---|---|
| HF- / SIS- Baseline vs Post SIS | 118.7 | 77.20 | 41.47 | 3.074 | 36 | * | 0.0364 |
| HF- / SIS+ Baseline vs Post SIS | 99.99 | 48.08 | 51.91 | 4.712 | 36 | ** | 0.0020 |
| HF+ / SIS- Baseline vs Post SIS | 107.3 | 64.32 | 42.95 | 3.183 | 36 | * | 0.0306 |
| HF+ / SIS+ Baseline vs Post SIS | 86.83 | 96.18 | -9.346 | 0.8484 | 36 | ns | 0.5523 |
| **Baseline** | | | | | | | |
| HF- / SIS- vs HF- / SIS+ | 118.7 | 99.99 | 18.68 | 1.175 | 72 | ns | 0.8396 |
| HF- / SIS- vs HF+ / SIS- | 118.7 | 107.3 | 11.40 | 0.6547 | 72 | ns | 0.9669 |
| HF- / SIS- vs HF+ / SIS+ | 118.7 | 86.83 | 31.84 | 2.003 | 72 | ns | 0.4935 |
| HF- / SIS+ vs HF+ / SIS- | 99.99 | 107.3 | -7.281 | 0.4580 | 72 | ns | 0.9882 |
| HF- / SIS+ vs HF+ / SIS+ | 99.99 | 86.83 | 13.16 | 0.9254 | 72 | ns | 0.9137 |
| HF+ / SIS- vs HF+ / SIS+ | 107.3 | 86.83 | 20.44 | 1.286 | 72 | ns | 0.8000 |
| **Post SIS** | | | | | | | |
| HF- / SIS- vs HF- / SIS+ | 77.20 | 48.08 | 29.12 | 1.832 | 72 | ns | 0.5689 |
| HF- / SIS- vs HF+ / SIS- | 77.20 | 64.32 | 12.88 | 0.7395 | 72 | ns | 0.9533 |
| HF- / SIS- vs HF+ / SIS+ | 77.20 | 96.18 | -18.98 | 1.194 | 72 | ns | 0.8332 |
| HF- / SIS+ vs HF+ / SIS- | 48.08 | 64.32 | -16.24 | 1.022 | 72 | ns | 0.8878 |
| HF- / SIS+ vs HF+ / SIS+ | 48.08 | 96.18 | -48.10 | 3.383 | 72 | ns | 0.0877 |
| HF+ / SIS- vs HF+ / SIS+ | 64.32 | 96.18 | -31.86 | 2.004 | 72 | ns | 0.4930 |
